# Analysis of the complete genome sequence for *Halococcus dombrowskii* ATCC BAA-364^T^

**DOI:** 10.1101/2022.08.16.504008

**Authors:** Sung W. Lim, Elizabeth G. Maurais, Ashlyn C. Farwell, Nicolette Barber, Abbey J. Olsen, Kristina F. Shalygina, Medina Omeragic, Eugenia A. Fedorov, Kyle S. MacLea

**Affiliations:** Binomica Labs, Long Island City, NY 11101 USA; Biotechnology Program, University of New Hampshire, Manchester, NH 03101 USA; Biology Program, University of New Hampshire, Manchester, NH 03101 USA; Neuropsychology Program, University of New Hampshire, Manchester, NH 03101 USA; Department of Life Sciences, University of New Hampshire, Manchester, NH 03101 USA

**Keywords:** Euryarchaeota, halobacteria, halococci, CRISPR, rRNA

## Abstract

We describe sequencing and assembly of complete *Halococcus dombrowskii* H4^T^ (=ATCC BAA-364^T^) genome using short- and long-read sequencing technologies. The first closed genome within its genus is composed of a 2,767,537 bp chromosome and five additional plasmids totalling 3,965,466 bp, with GC content of 62.18%. The genome contains 4,029 genes, 3,963 coding sequences and two CRISPR arrays. Unusually, this Euryarchaeote carries multiple rRNA operons with divergent ITS identities across both its chromosome and plasmids.

## INTRODUCTION

Among the halophilic archaea, the order Halobacteriales is notable for containing a range of organisms with extreme tolerance for salt, including the most halophilic organisms known (1). These archaea are found abundantly in salterns, salt flats, and other salt-rich environments, and are frequently accompanied by pink or red pigmentation, documented as far back as very old Chinese literature (1). Since the earliest characterization of the halobacteria in the modern era, these diverse salt-loving bacteria have been a source of interest to biologists.

A range of similar halobacteria observed in the early days of microbiology were often not well described, which led to problems of taxonomic circumscription in an era when genetic and genomic characterization was not yet possible. Members of the halobacterial genus *Halococcus* were no exception, with a likely early observation in pigment-induced reddening of flesh in the New England cod fisheries of 1879 (2). Later, the taxonomy of *Halococcus—*as being distinct from other Halobacteriales—was firmed up, with valid publication of the genus in 1935 (3–5). Despite early publication of the genus, collection of a few distinctly-reported strains into a single species in that genus (*Halococcus morrhuae*, the type species) did not happen until 1973 (4).

This first member of the genus, *Halococcus morrhuae*, is recognized to have been identified initially during the examination of the reddened codfish from 1879 (2, 4, 6). Expansion of the genus with the discovery of the other nine validly-published members did not occur until much later between 1990 and 2018 (7–15). Salt-containing natural environments and commercial salt products remain the dominant sources for identified halococci, although they have also been found in the leaf tissue of Cabo Rojo mangroves (16), in the nostrils of seabirds (17), and in Thai fish sauce (12).

Eight of the validly published *Halococcus* species have genome drafts available, along with one additional environmental isolate (16, 18–21); none of the available genome sequences are complete.

One of the unsequenced species is *Halococcus dombrowskii. H. dombrowskii* is particularly notable for its isolation from sections of Permian subsurface salt deposits (9) estimated to date back as far as 225 to 280 million years ago (8, 22). The question of whether the halococci found in ancient salt deposits could be considered molecular fossils of the distant past, signs of isolated ecosystems, or even relatively modern contamination remains unresolved, but it does beg further investigation (23).

To aid in understanding the evolutionary relationships within the *Halococcus* genus, as well as to fill the void in sequenced genomes in this group of organisms, we sequenced *Halococcus dombrowskii* H4^T^ (=ATCC BAA-364^T^). As detailed below, we were able, using long-read sequencing technology (polished with higher quality short-read data), to close the main chromosome as well as the sequences of 5 additional *Halococcus* plasmids. These sequences provide useful information with which to compare and study other *Halococcus* genomes and may help answer other questions in the field of halobacteria more broadly.

## METHODS

### DNA isolation and sequencing of *H. dombrowskii* ATCC BAA-364 ^T^

Isolation of genomic DNA and sequencing of *H. dombrowskii* ATCC BAA-364^T^ was undertaken directly from double-walled glass vials purchased from ATCC (Manassas, Virginia, USA). Separate glass vials were used for the short-read library prep and for each of two long-read library preps (detailed below).

For short-read sequencing, genomic DNA (gDNA) was isolated from half of the lyophilized cell pellet using the FastDNA SPIN Kit for Soil using the standard protocol and lysing matrix A—a mixture of garnet and ceramic—on a FastPrep 24 homogenizer. The homogenizer, kit, and matrix were purchased from MP Biochemicals (Irvine, CA, USA). The sequencing library was generated from the gDNA input using the KAPA HyperPlus kit (KR1145, v.5.19, kit KK8515, Wilmington, MA, USA) by enzymatic fragmentation with HyperPlus end repair. The Hubbard Center for Genome Studies at the University of New Hampshire (HCGS, Durham, NH, USA) ran this library on an Illumina HiSeq 2500 sequencer, producing 250-bp paired-end fragments.

For long-read sequencing, two separate genomic DNA isolations and library preps were undertaken. For the first long-read sample, designated M2, a different sample from the short-read sample (in its own double-walled glass vial), containing a lyophilized cell pellet of ATCC BAA-364, was opened and the pellet was rehydrated in 400 μl of 1x phosphate-buffered saline (PBS). After spinning down (at 16,000 x g) 200 μl of the well-mixed sample, the supernatant was removed. 20 μl of 1x PBS was added to the cell pellet and cells were mixed 10x with a P200 pipet with a standard tip. 100 μl of TESTS-LYS buffer (50 mM Tris-HCl, pH 8.0, 50 mM EDTA, 8% (w/v) sucrose, 5% (v/v) Triton X-100, 0.5% (v/v) SDS, 10 mg/ml lysozyme) was added to the cells and PBS mixture, followed by vortexing 10 × 1s on maximum power. The sample was incubated at 37°C for 75 minutes and then employed the Circulomics Nanobind CBB Big DNA Kit (NB-900-001-01, Baltimore, MD, USA). The CBB kit used the Circulomics EXT-GPH-001 protocol beginning at step 5, the addition of proteinase K, and proceeded as directed. After the DNA rested overnight at room temperature per the kit instructions, the DNA was mixed 10x using a P200 pipet with a standard tip to produce the purified gDNA.

M2 genomic DNA was used as input to the ligation sequencing kit LSK109 (Oxford Nanopore Technologies (ONT), Oxford, UK). The resulting M2 Nanopore library was run on a MinION R9.4.1 flowcell (FLO-MIN-106) on an ONT GridION instrument (at HCGS) running MinKNOW 21.05.20 (MinKNOW Core 4.3.11). Sequences were basecalled using guppy basecaller v.5.0.13 (24) in SUP mode.

A second genomic DNA isolation and library prep was undertaken for long-read sequencing. This procedure was identical to the M2 process above with the following exceptions: The entire 400 μl rehydrated cell pellet from a new double-walled glass vial of ATCC BAA-364 was employed for sample M3, versus the 200 μl (half pellet) used in M2. The initial 37°C incubation for M3 was two hours in length. All other steps for M3 were identical to the steps used in M2 above.

### Software commands for genome analysis and assembly

Processing of Nanopore MinION (long) reads and Illumina (short) reads as well as analysis, assembly, and characterization of the subsequent genomes, different software programs were carried out on a system running Pop! OS 20.10 64bit with Ryzen 9 5950x and 64GB ECC RAM. Programs and pipelines used for the process are described in summarized form below. In each case, detailed arguments and options used for each of the specific software commands are found in the Supplementary Information for this manuscript, in the Supplementary Methods section.

### Characterizing and filtering raw MinION and Illumina reads

All MinION reads from the M2 and M3 samples were characterized using NanoPlot v.1.39.0 (25). We then used FiltLong v.0.2.1 on only the pass-quality reads from M2 and M3 (minimum Q score of 10) to generate working data for each group, screening for minimum read length of 1000 bp and then keeping the top scoring 90% of the output from each group. The resulting working read sets were characterized with NanoPlot a second time. We observed fragmented (high contig numbers) Flye-based (26) exploratory assemblies based solely on M2/M3 data. To address the possibility that we were working with highly-degraded DNA, we also combined the M2 and M3 passed reads into a single set prior to the filtering process described above. The combined reads were filtered separately as a third data group referred to as M23.

For use in polishing subsequent assemblies, we also filtered the short-read Illumina data (referred to as HdomA3) using fastp v.0.23.1 (27).

### Genome assembly

Trycycler v.0.5.3 (28) provided the backend for *Halococccus dombrowskii* genome assembly, based on output of four different assemblers; Flye v.2.9-b1768 (26), Miniasm v.0.3-r179 (29) & Minipolish v0.1.3 (30), Raven v.1.7.0 (31), and finally Shasta v.0.8.0 (32). M2, M3 and M23 filtered read sets were used to create 16 subsampled reads each. The subsampled reads corresponded to 4 assemblies per 4 different target assemblers, producing 48 assemblies in total.

A bash script was used to generate assemblies from each of the subsampled reads (Script available from https://github.com/naturepoker/Halococcus_WIP_scripts), and resulting assemblies were annotated with Prokka v.1.14.6 (33) to check for rRNA and CRISPR spacer site counts (utilized in subsequent analysis). GToTree v.1.6.31 (34, 35) was used to generate cladograms for each of the M2/M3/M23 set-based assembly groups, and the annotation file was curated manually to show rRNA and CRISPR count per assembly using iTOL v5 (36) on the <u>itol.embl.de</u> visualization platform.

18 assemblies were chosen out of 48 based on rRNA count and CRISPR spacer site count (detailed in Results and Discussion) and pooled into a separate directory to create clusters of raw reads around the contigs. We used the filtered M23 reads for this step. The clustering process generated 39 clusters containing 165 contigs in total. Reconciliation of the contigs using the Trycycler (28) reconcile option filtered the clusters down to a final 6, clusters 1, 2, 3, 4, 6 and 9. Note that the reconciliation command was run on a per-cluster basis. Multiple sequence alignment (msa) was performed on each of the remaining clusters using Trycycler’s msa option. The filtered M23 reads were partitioned to each of the aligned cluster sequences, final consensus sequence was generated on a per contig basis, and the contigs were combined into the final raw assembly.

The raw assembly was polished using an in-house script (Script available from https://github.com/naturepoker/Halococcus_WIP_scripts) using bwa v.0.7.17-r1198-dirty (37), Racon v.1.4.20 (38), <u>Medaka</u> v.1.5.0, and Polypolish v.0.5.0 (39).

### Characterizing and annotating the genome

We performed additional QC to assess completeness of the genome in terms of essential genes by benchmarking universal single-copy orthologs (BUSCO) analysis, with v.5.3.2 (40–42) and the archaea_odb10 profile.

GToTree (34) was used again to generate a whole genome phylogenetic tree of *Halococcus dombrowskii* against other *Halococcus* spp. genomes in NCBI. *Halococccus* reference genomes were queried on Genbank, returning 10 genome accession numbers, for *Halococcus hamelinensis* 100A6 (sequenced twice), *Halococcus saccharolyticus* DSM 5350, *Halococcus salifodinae* DSM 8989, *Halococcus morrhuae* DSM 1307, *Halococcus thailandensis* JCM 13552, *Halococcus salsus* ZJ1, *Halococcus sediminicola* CBA1101, *Halococcus agarilyticus* 197A, and *Halococcus* sp. IIIV-5B.

In preparation for a more detailed Prokka annotation, we curated a list of known genomes from class Halobacteria on NCBI—returning 501 genome accessions—using <u>NCBI-genome-download</u> v.0.3.1. The protein multifasta files from these 501 genome accessions were used to create a halobacterial protein sequence file. To annotate and characterize our genome assembly we annotated each contig separately using Prokka (33) and the halobacterial protein sequence library derived from the 501 accessions above.

### Visualization and analysis *Halococcus* spp. genomes

A docker instance of CGview Comparison Tool (43) was used to compare *Halococcus dombrowskii* against other *Halococcus* spp. at the whole genome level. NCBI-genome-download v.0.3.1 was used again to obtain Genbank files of the 10 available *Halococcus* spp. genomes to enable the CGView comparison. rRNA regions and downstream genes were manually isolated using samtools v.1.14 (44) with both Prokka-generated and NCBI Genbank files as reference. The DNA sequences around 10,000 bases in length were re-annotated with Prokka and used with <u>Geneblocks</u> v.1.2.2 script (Script available from https://github.com/naturepoker/Halococcus_WIP_scripts) to generate figures used in this paper. The same process was used to isolate the Internal Transcribed Spacer (ITS) sequences from the *Halococcus* genomes, and Clustal Omega (45), hosted on the website of the <u>EBI</u>, was used to align them and generate the cladogram used for the figure in this paper. (Full size figures are available on https://github.com/naturepoker/Halococcus_WIP_scripts)

### Isolating reads unique to *H. dombrowskii* rRNA regions

M23 raw reads were filtered again with a minimum length of 10,000 bp to cut down on spurious mapping to only parts of our regions of interest. The top 90% of reads of at least 10,000 bp in length were isolated using <u>filtlong</u> v.0.2.1 (R.R. Wick).

Three sequence blocks containing rRNA operon and downstream genes were manually isolated from the *Halococcus dombrowskii* genome with samtools (44) and Prokka-annotated genbank file: the 11150 bp chromosomal (contig 001) region (NZ_CP095005.1) between 209244-220393, the 9117 bp plasmid2 (contig 003) region (NZ_CP095007.1) between 126264-135380, and the 10433bp plasmid4 (contig 006) region (NZ_CP095009.1) between 168572-179004.

Minimap v.2.24-r1122 (29) and Bedtools v.2.29.1 (46) were used to isolate reads mapping uniquely to the rRNA operon and downstream gene regions on either the chromosomal rRNA, plasmid2 rRNA copy, or plasmid4 rRNA copy, in our *H. dombrowskii* genome. We used the linux grep command to compare and isolate unique fastq headers from reads mapped to each of the sequence blocks. For each sequence block (main chromosome, plasmid 2, plasmid 4), the unique read headers were sorted and combined to create a list we could use to compare against reads aligned to the other sequence block(s). <u>Seqtk</u> v.1.3-r117-dirty was then used to create fastq files containing only reads unique to each of the sequence blocks. NanoPlot (25) visualized the unique reads fastq files to allow characterization of the distinct sequence blocks. The Integrated Genomics Viewer (IGV) (47) was also used to show mapped reads against the rRNA operon and downstream gene regions.

## RESULTS AND DISCUSSION

### Exploratory genome assemblies based on raw long and short reads

We started our genome assembly with the Guppy-basecalled reads (SUP mode) from 2 MinION flowcells (M2 and M3), along with one paired-end short read set from Illumina HiSeq. M2 data consisted of 378,911 reads totaling 2,359,879,393 bases with N_50_ of 17,233 bases. M3 data consisted of 2,363,742 reads totaling 9,544,901,750 bases with N_50_ of 21,871. Illumina data consisted of 4,327,290 reads totaling 1,046,384,000 bases.

Initial Flye-based (26) exploratory assemblies showed a high degree of size, circularization, and contig number variations despite close to 1500× read depth across the available reads Subsampling the raw reads to a more reasonable 200-300× coverage or applying more stringent quality score and/or minimum read length cutoffs did not result in more uniform contiguity. One exception to the observed fluctuation was the circularized main chromosome carrying an RNA operon in tRNA-Cys, 5S, 23S, tRNA-Ala and 16S configuration along with a CRISPR spacer site containing 118 and 13 repeat units. We found a varying number of additional rRNA operons and CRISPR spacer sites always located on non-chromosomal contigs as well.

### Trycycler multi-assembly pipeline

Assembly fluctuations from the exploratory assemblies prompted us to utilize the Trycycler (28) pipeline on minimally-filtered (above Q10, above 1kb, top 90% scoring) reads from M2 or M3 MinION data and a combined set of both (referred to as M23). The decision to test out a combined read set of M2 and M3 as a third separate dataset was made based on a possibility our *Halococcus dombrowskii* DNA had been exposed to genome shearing damage or higher than expected transposition activities prior to the extraction and sequencing process. The newly filtered M2 data consisted of 134,800 reads totaling 1,259,984,769 bases with an N_50_ of 19,232 (Fig. 1A). New filtered M3 data consisted of 446,376 reads totaling 4,803,540,899 bases with N_50_ of 25,476 (Fig. 1B). M23, which is a combination of M2 and M3 data filtered with the above criteria, consisted of 579,763 reads totaling 6,063,484,969 bases with N_50_ of 24,231 (Fig. 1C).

**Figure 1.**
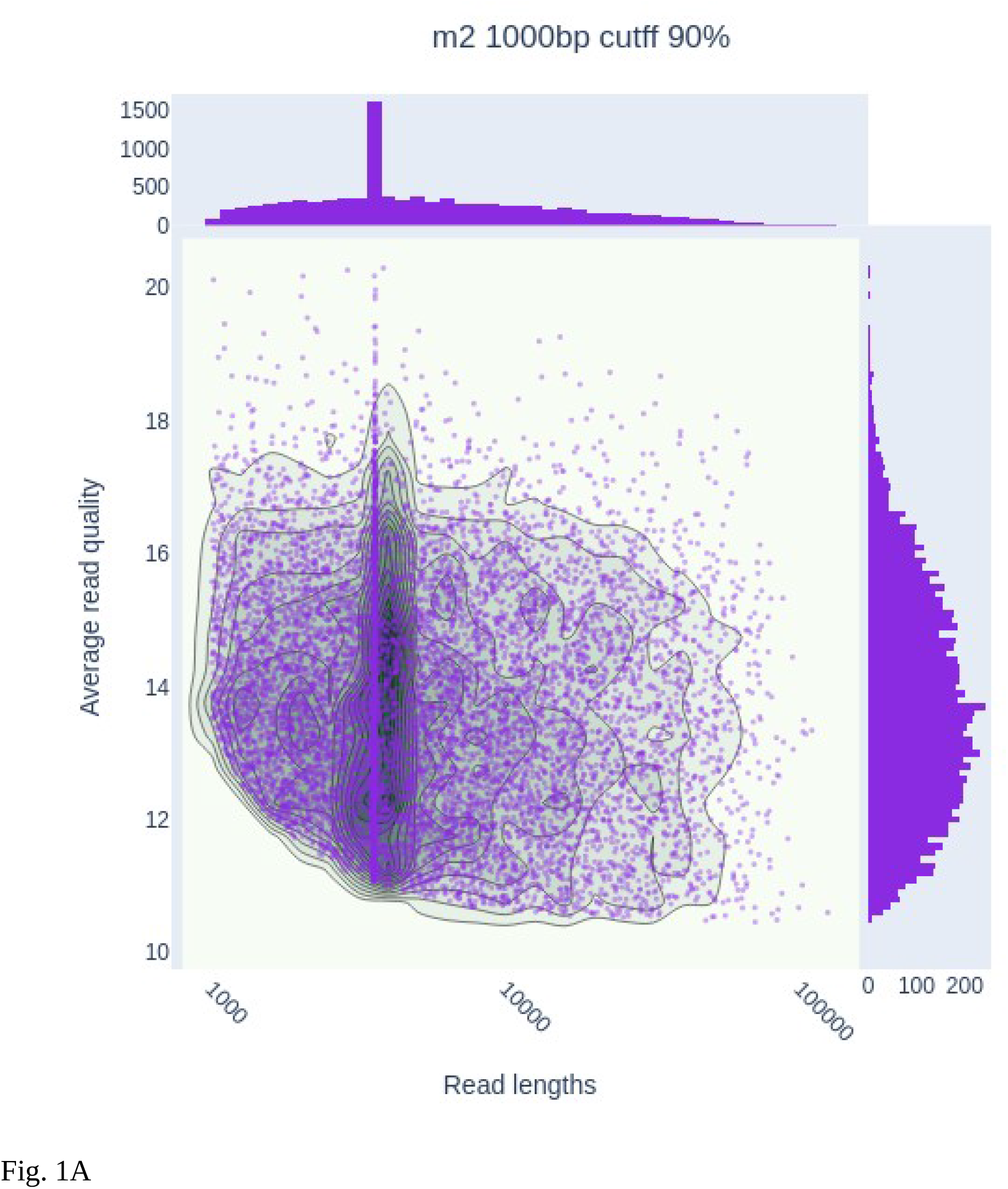
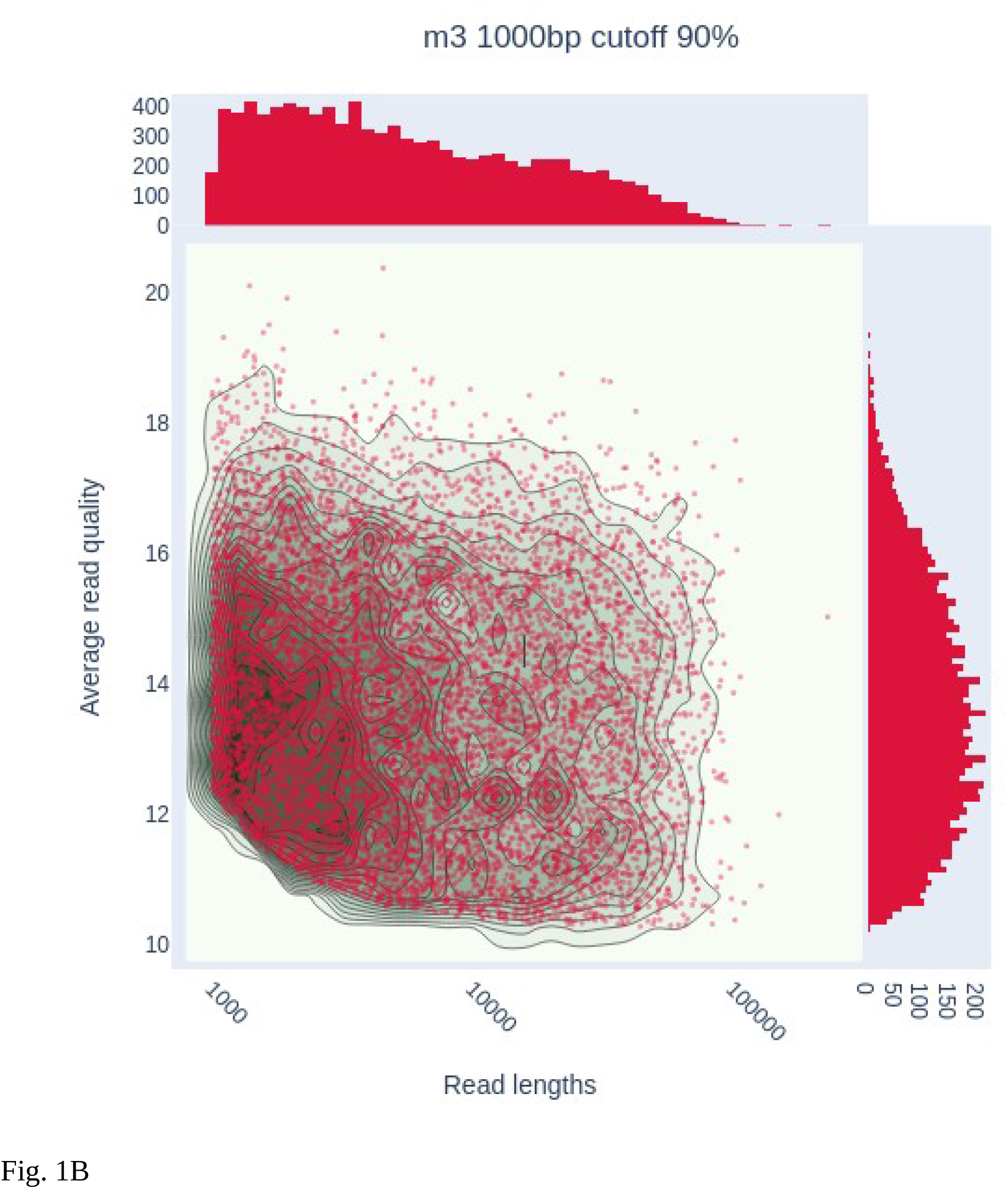
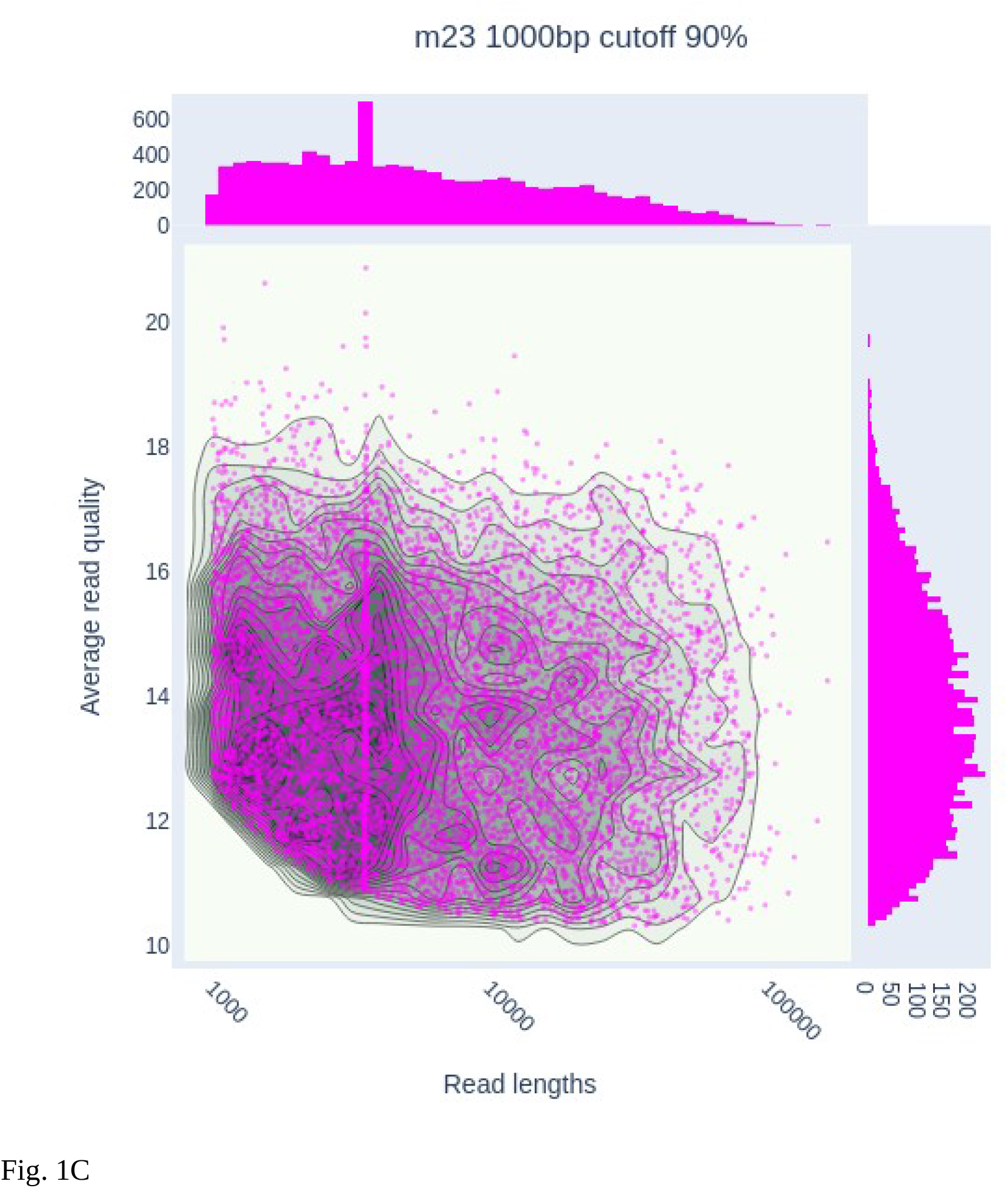
NanoPlot output of the three working long-read sets, filtered for top 90% reads by read length and average quality score among reads meeting minimum threshold of q10 average score and at least 1000 bp in length.

The three read sets were subsampled into 16 groups each, for a total of 48 subsampled read sets. Each of the subsampled reads were assigned to one of four assemblers (Flye (26), Miniasm (29), Raven (31), Shasta (32)), creating 16 assemblies per set, 48 assemblies in total. Annotation of the new assemblies agreed with our previous observation of non-chromosomally located rRNA operon sites across all 48 of them. The number of the operons ranged from two to five with independent rRNA gene counts ranging from 6 to 15. All of the rRNA operons were located on separate, independent contigs. Of the 48 assemblies, 24 showed the dominant rRNA operon distribution pattern consisting of three rRNA operons containing 16s, 23s and 5s, making for a total of 9 rRNA sites on the genomes, spread among three separate contigs.

Distribution of CRISPR spacers in the 48 assemblies showed a dominant pattern as well. There were only two different types of CRISPR spacers across all 48 assemblies (Fig. 2A-C). Chromosomal CRISPR spacer sites consisted of 118 and 13 repeat units, and another pattern showed up on non-chromosomal contigs coding for rRNA operons consisting of 8, 34, and 38 repeat units. All instances of multiple non-chromosomal CRISPR spacer sites turned out to be replicates of the 8,34,38 repeat unit pattern, leading us to suspect they were duplications. Out of 48 CRISPR spacer sites annotated across the assemblies, 38 showed the pattern of two spacer sites per genome, containing the repeat units described above.

**Figure 2.**
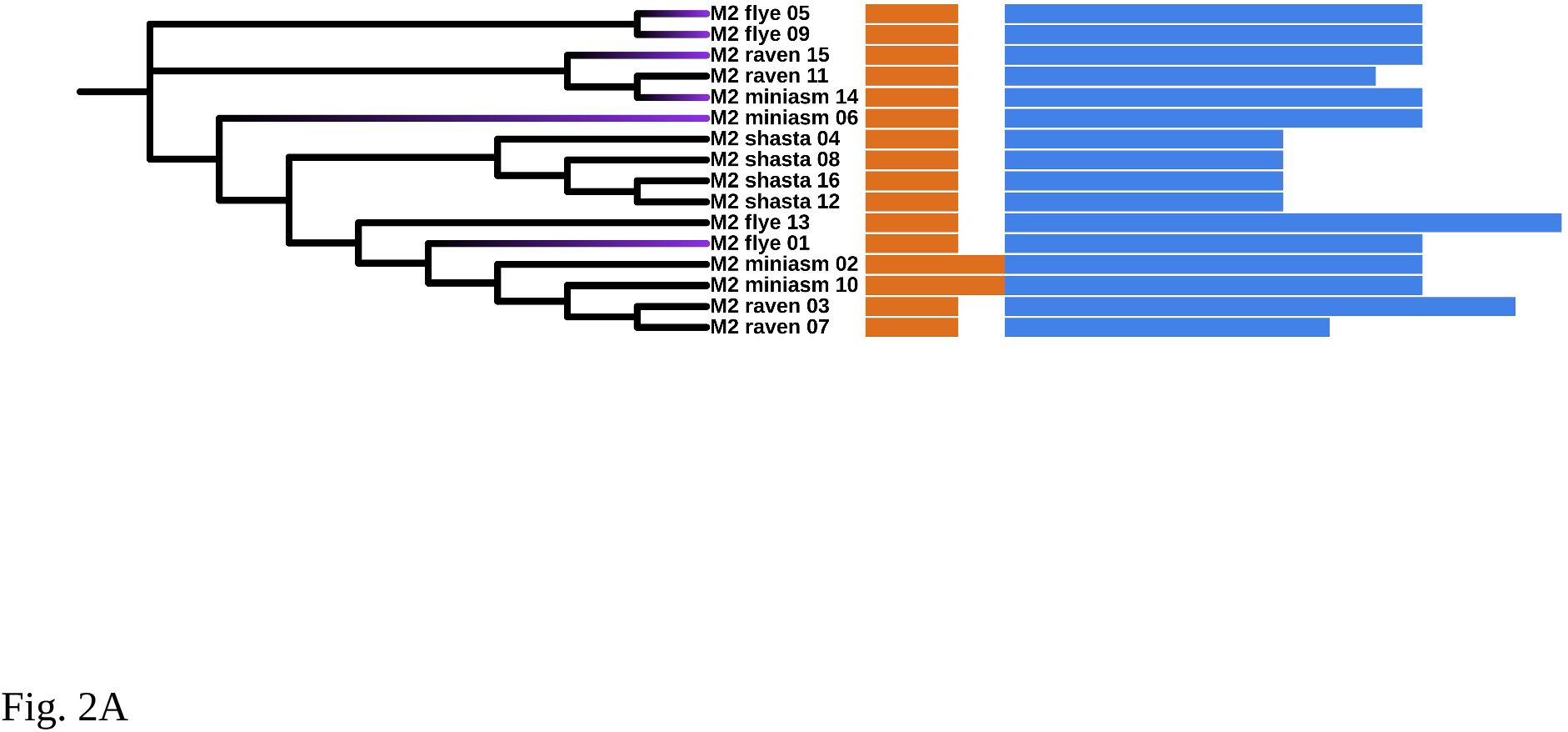
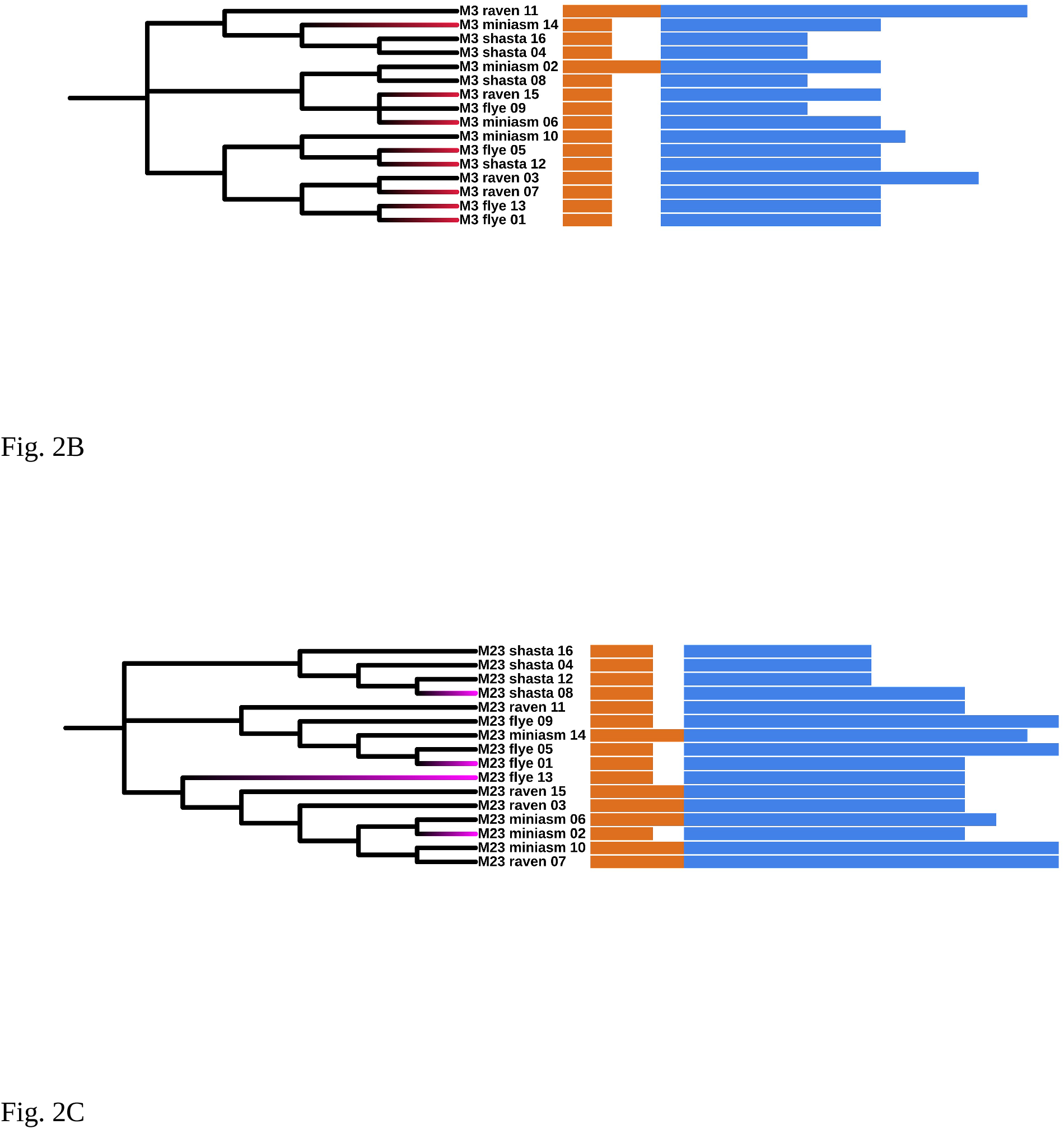
Cladogram of subsampled assemblies from group M2, M3 and M23 built with GToTree’s (34) Archaeal SCG set. The orange bar shows the CRISPR spacer array counts, ranging from 2 to 4. The blue bar shows individual rRNA counts per assembly, ranging from 6 to 15. Assemblies chosen for final consensus generation are highlighted in color.

Based on the rRNA operon and CRISPR spacer site distribution pattern, we sorted out only the assemblies carrying three rRNA operons, where one operon must be paired with a CRISPR spacer site of 118 and 13 repeat units on a chromosomal contig, and other operon must be paired with a CRISPR spacer site of 8, 34, and 38 repeat units on a non-chromosomal contig.

This left us with a curated set of 18 assemblies consisting of 6 assemblies from M2 flowcell, 8 assemblies from M3 flowcell, and 4 assemblies from the M23 combined read set (Fig. 3). The final group of assemblies represent 8 Flye outputs, 5 Miniasm outputs, 3 Raven outputs and 2 Shasta outputs. The manually-curated assembly set was then processed through the conventional Trycycler pipeline and polished with a round of Racon (38), Medaka, and Polypolish (39), the last step using the *H. dombrowskii* short-reads obtained from the Illumina HiSeq platform.

**Figure 3.**
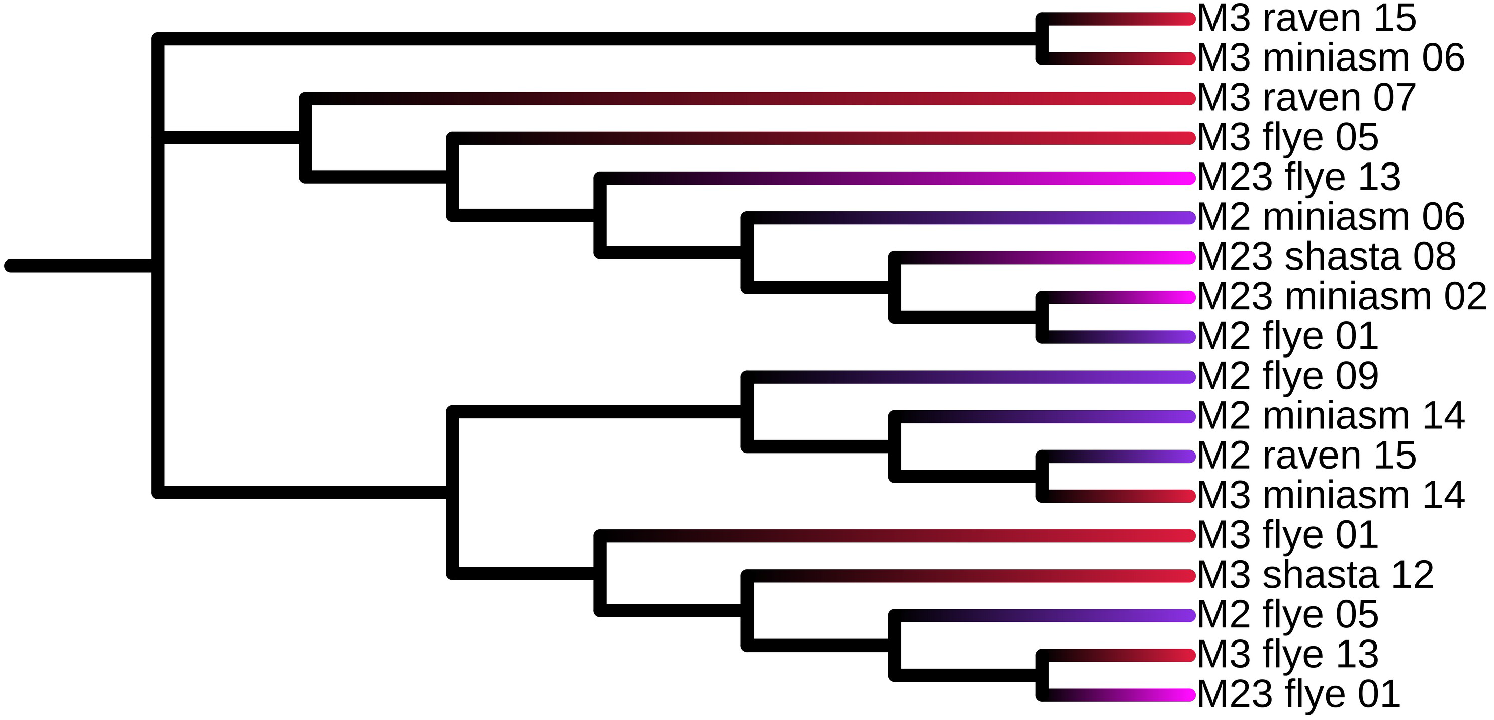
Cladogram of the final 18 assemblies chosen from the initial 48 assemblies based on rRNA and CRISPR spacer array counts, based on GToTree’s Archaeal single-copy gene (SCG) set.

Screening of the polished genome by benchmarking universal single-copy orthologs (BUSCO) analysis (40–42) with the archaea_odb10 dataset shows 99.5% recovery of essential genes without duplication or fragmentation, with the polished genome missing only a putative diphthine synthase ortholog from the 194 BUSCO gene set.

### Genome overview and comparison

The *Halococcus dombrowskii* genome (<u>GCF_022870485.1</u>) is 3,965,466 bp long with GC content of 62.18%, coding for 4029 genes and 3963 coding sequences (CDS) across six circularized contigs (Table 1). The main chromosome accounts for ∼69.8% of the total size of the genome, which is consistent with earlier estimates that *Halococcus morrhuae* (the type species of the genus) is made up 70% non-satellite DNA and 30% satellite DNA (representative of plasmid DNA) (1).

**Table 1.**
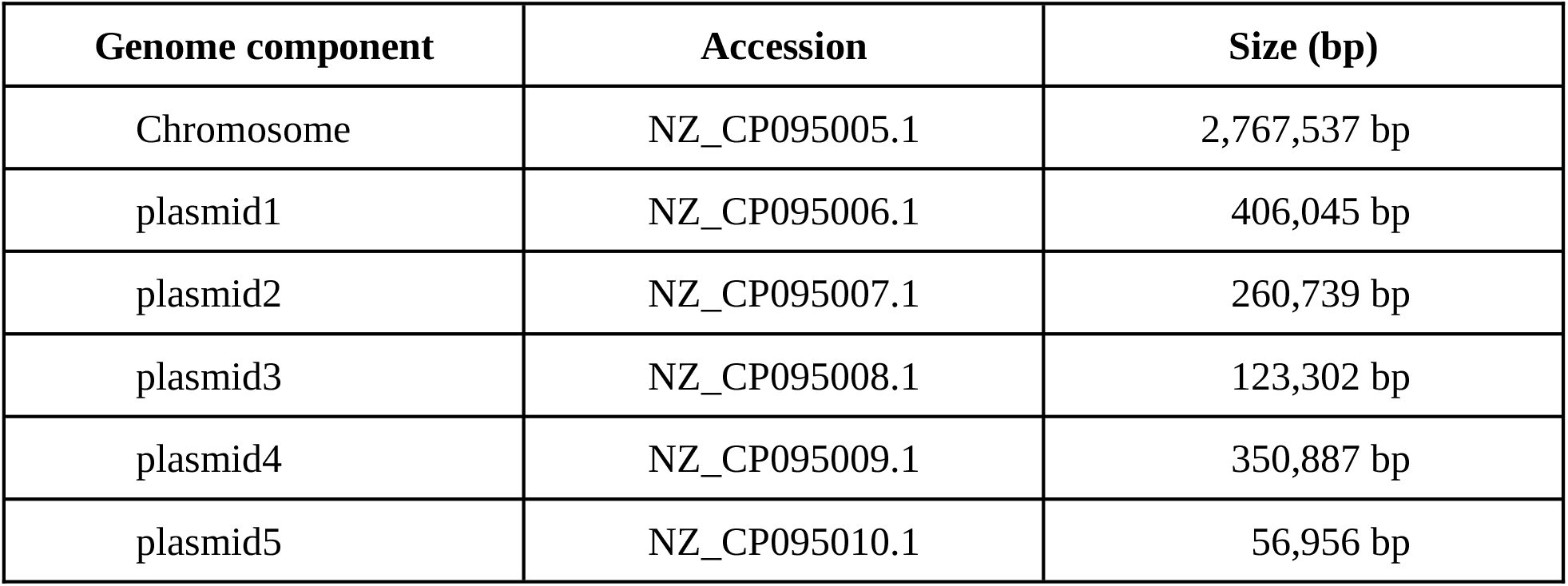
*Halococcus dombrowskii* genome components with NCBI RefSeq accessions and sizes of each component indicated (in bases).

The genome contains 9 rRNA genes, 55 tRNAs and 5 CRISPR arrays across its chromosome and plasmids. The 9 rRNA sites are split among the chromosome, plasmid2 and plasmid4 as three polycistronic operons containing tRNA-Ala in their intergenic spacer regions. The CRISPR sites are split between the chromosome and plasmid4, with the chromosome carrying two spacer sites of 118 and 13 repeat units, and plasmid4 carrying three spacer sites of 34, 38, and 8 repeat units.

Comparing the *H. dombrowskii* genome against those of other known *Halococcus* species suggests a highly varied genomic landscape among the members of the genus, detecting significant synteny only between *H. dombrowskii* and its phylogenetically closest cousins, *H. thailandensis* (<u>GCF 000336715.1</u>) and *H. morrhuae* (<u>GCF 000336695.1</u>). Even that level of similarity drops off dramatically when comparing extrachromosomal contigs against other known *Halococcus* genomes (Fig. 4). This genomic variation seems to follow the pattern of evolutionary distance from one *Halococcus* organism to the others, as the degrees of sequence similarity on the chromosomal level closely follows phylogeny of the 11 *Halococcus* genomes based on 76 archaeal SCGs (Fig. 5).

**Figure 4.**
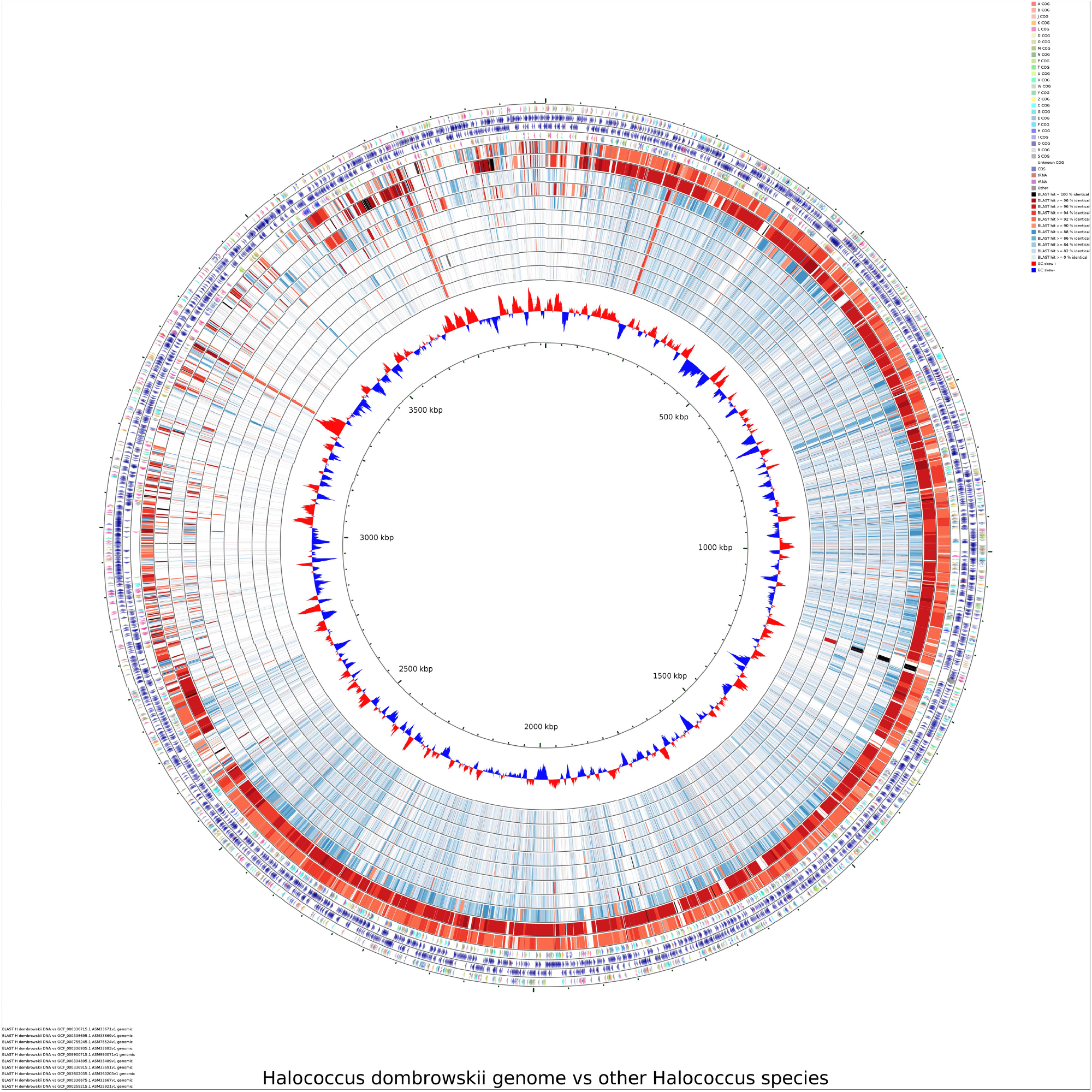
Whole genome comparison of the *H. dombrowskii* genome against other NCBI *Halococcus* spp. reference genomes. Outermost ring shows our assembly along with CDS and color-coded COG (Cluster of Orthologous Groups) (48) matches. Inner rings show other *Halococcus* genomes in the order of similarity to *H. dombrowskii*.

**Figure 5.**
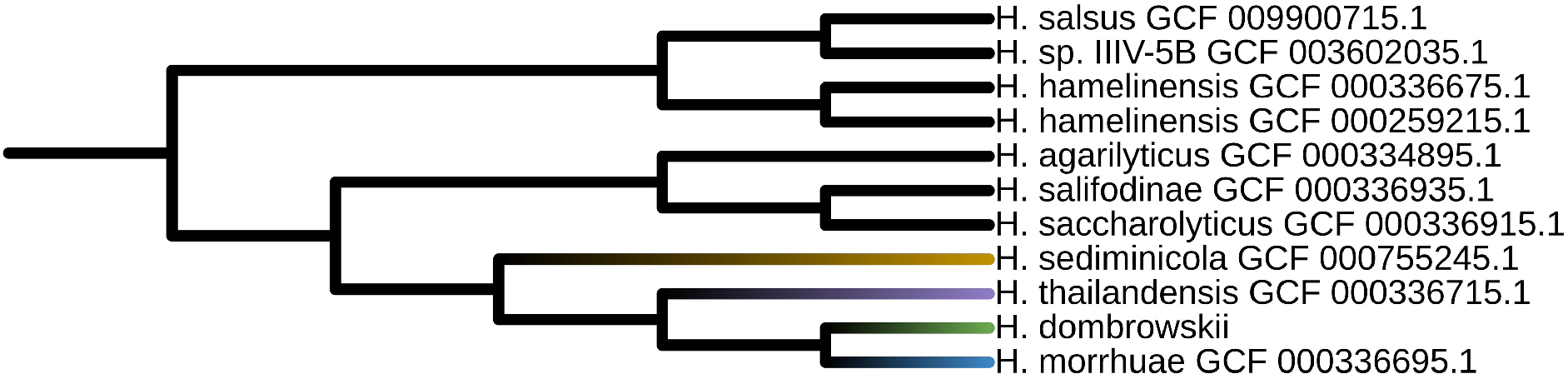
Whole genome phylogenetic tree of *Halococcus* species using 76 Archaeal SCGs, midpoint rooted. The clade closest to the *H. dombrowskii* genome is color-coded. The phylogenetic distribution closely follows the arrangement seen in the CCT full genome DNA to DNA comparison.

It should be noted that despite significant variations seen on the sequence level, *Halococcus* chromosomes still maintain a high degree of similarity on CDS protein sequence level (Fig. 6). The degrees of variation among the *Halococcus* CDS largely follows that of the SCG based phylogenetic tree as well, as demonstrated in the below comparison. It should be noted that just as with pure DNA sequence level comparison, the degree of homology among protein CDS drops off dramatically when we compare extrachromosomal regions against each other.

**Figure 6.**
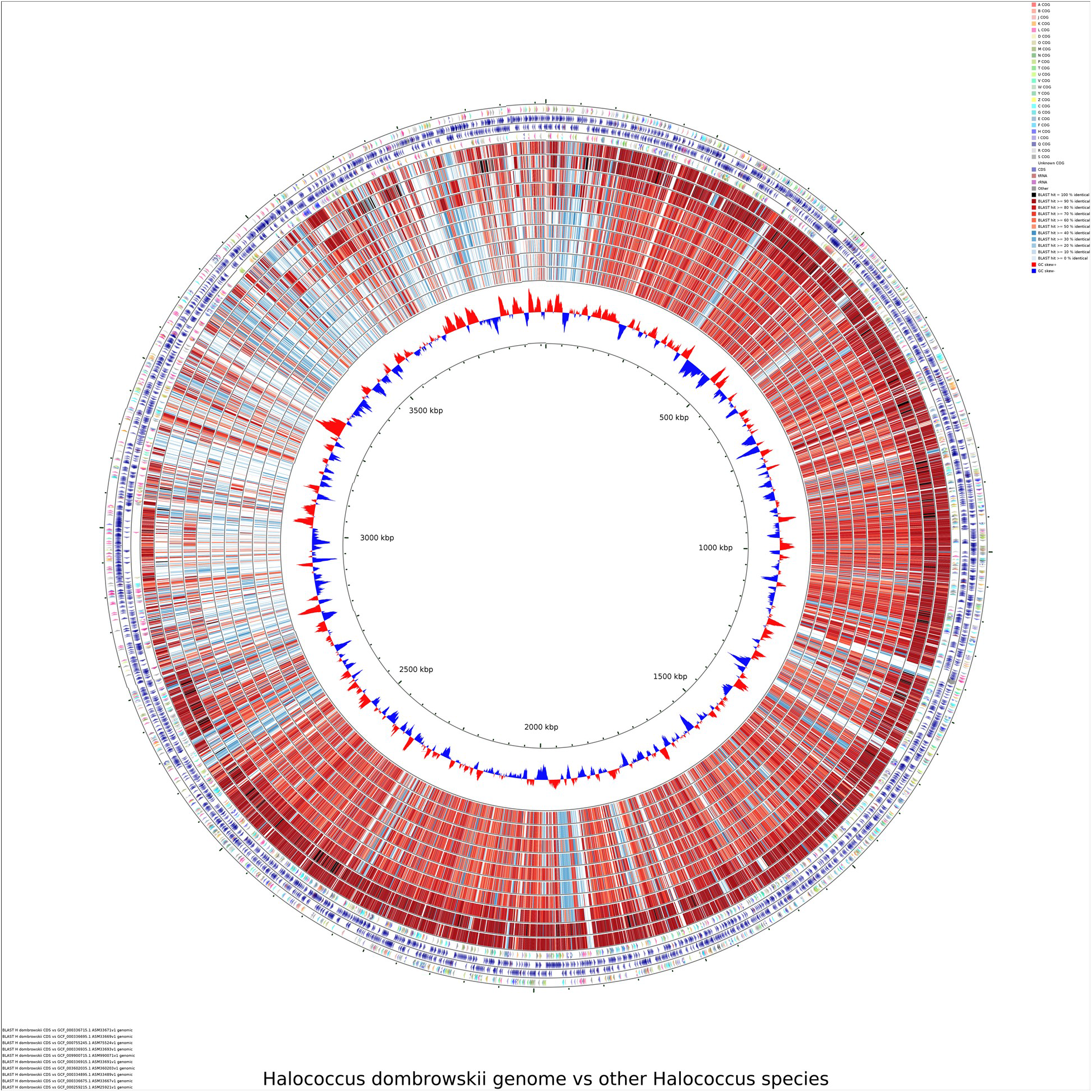
*Halococcus* genus CDS-to-CDS comparison created with CCT. The genomes are arranged in the order of similarity from the outermost ring (closest to *H. dombrowskii*), to the innermost ring, with redder color indicating higher CDS to CDS similarity. Starting from the top and running clockwise, we can see the similarity among even closely related genomes drop off as we compare the extrachromosomal regions of *H. dombrowskii* genome against other members of the genus.

### rRNA operon characterization

As of the time of this writing, *Halococcus dombrowskii* represents the first genome assembly in the genus with more than one observed rRNA operon. The chromosomal rRNA operon is oriented in typical 5S, 23S, 16S fashion along with tRNA-Cys and tRNA-Ala, much like previously studied examples of Euryarchaeota such as *Halobacterium cutirubrum* (49). All of the 11 *Halococcus* genomes surveyed in our study show a conserved series of genes further downstream of the rRNA operon of potential interest (Fig. 8A-8K), as shown below.

**Figure 7.**
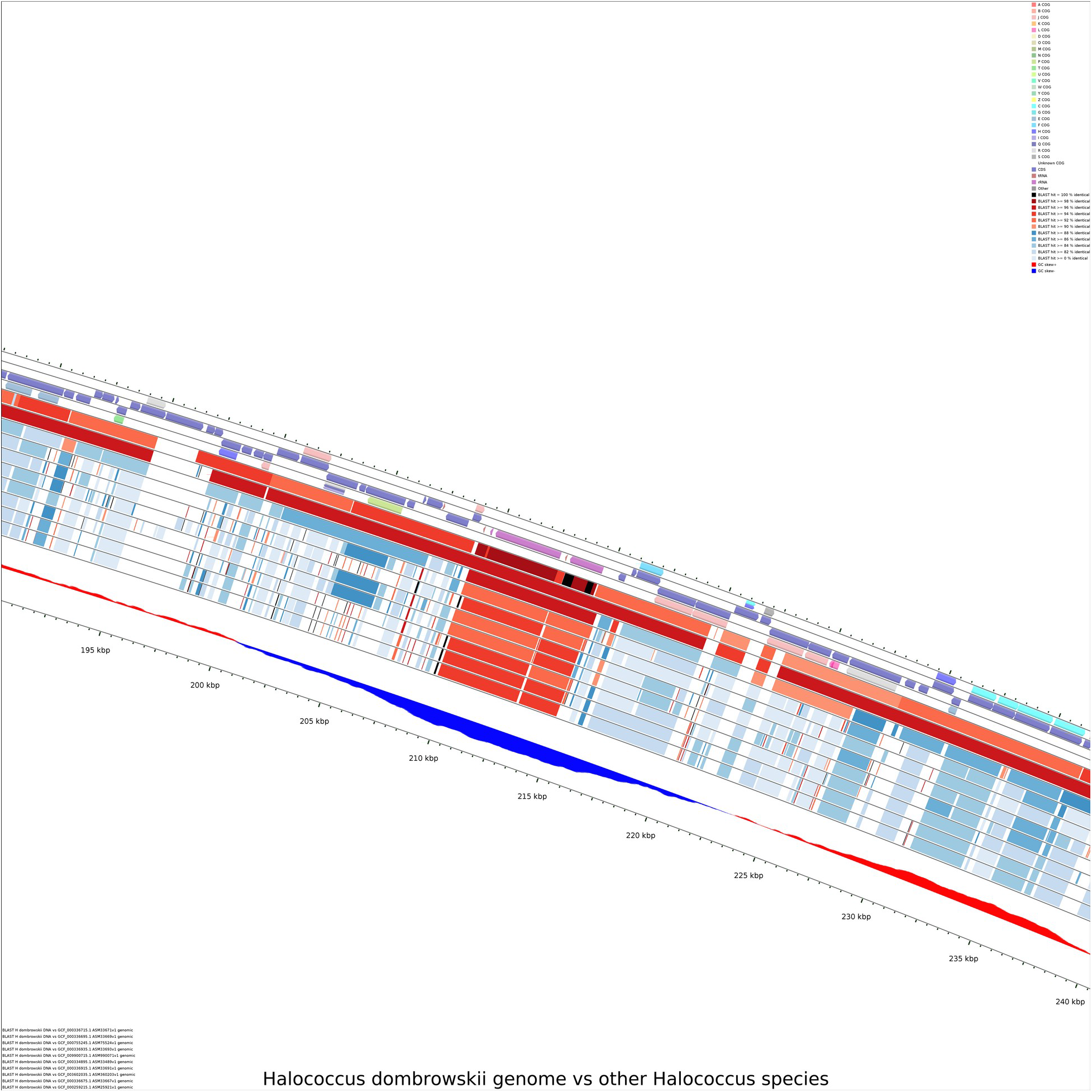
The region coding for rRNA operon (indicated with pink bar on the outermost *H. dombrowskii* genome track) displays conserved rRNA operon location - note that the second line below *H. dombrowskii* track showing *H. thailandensis* contains significant gaps carried over from the contig-level assembly, indicated with black bars.

**Figure 8.**
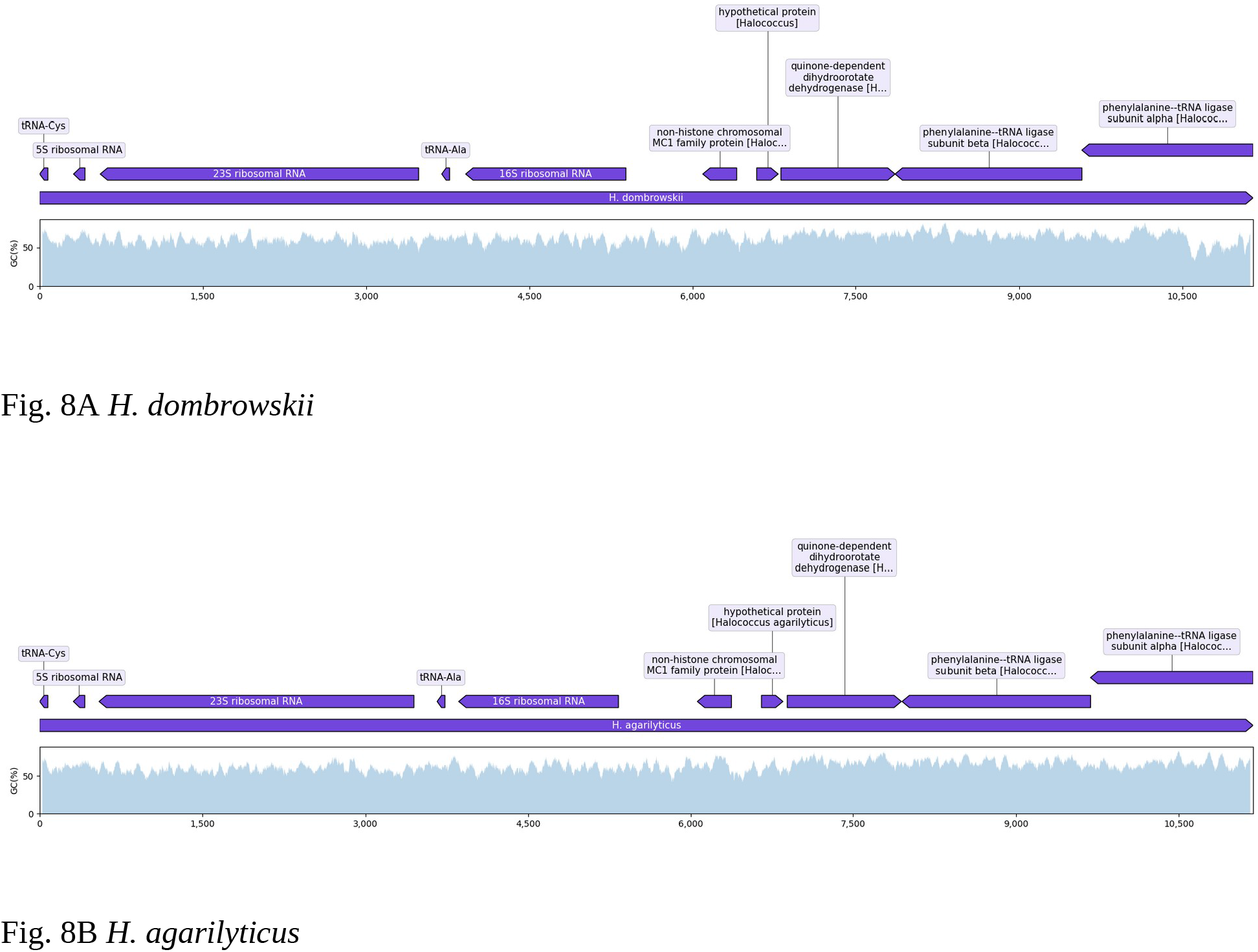
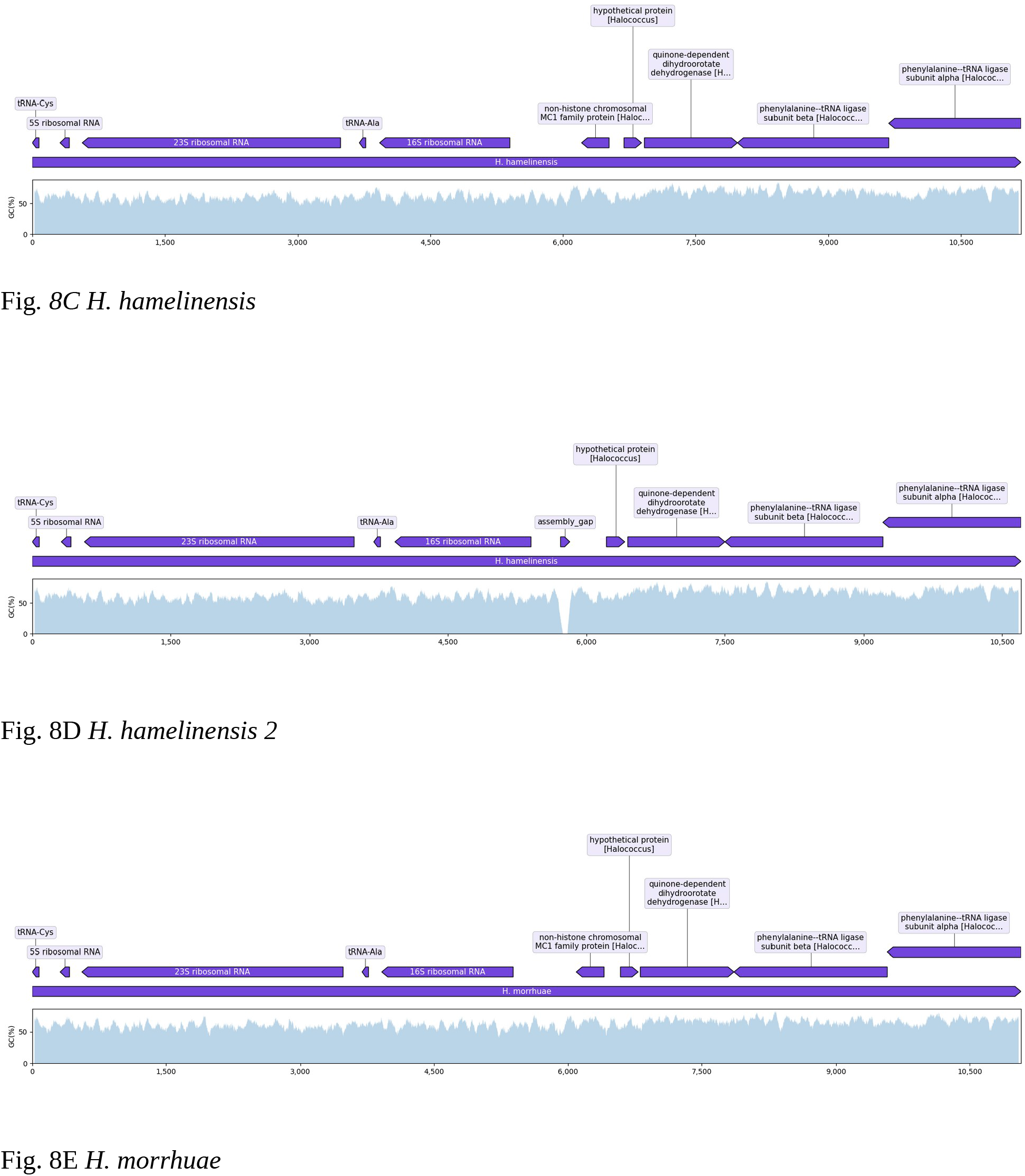
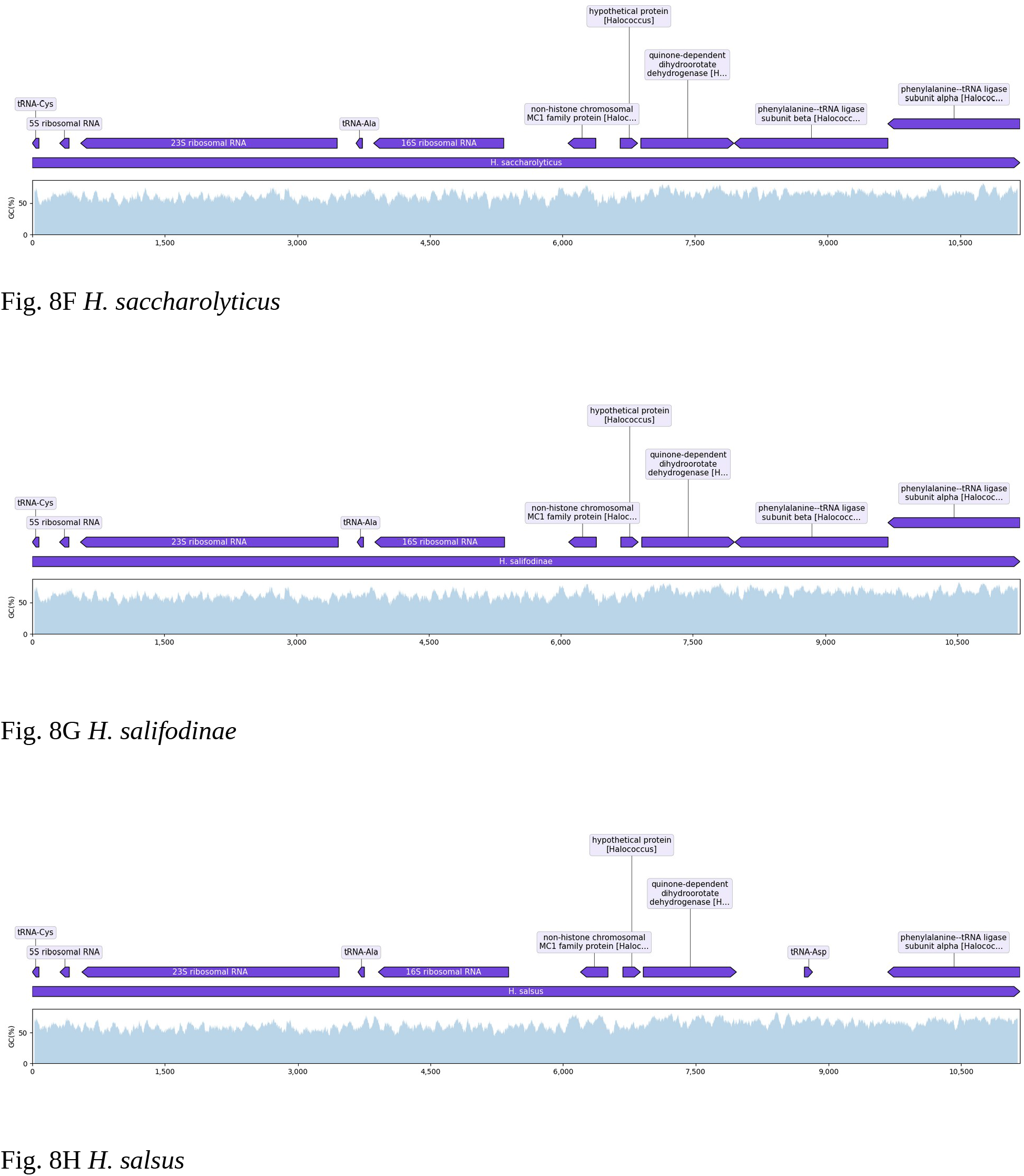
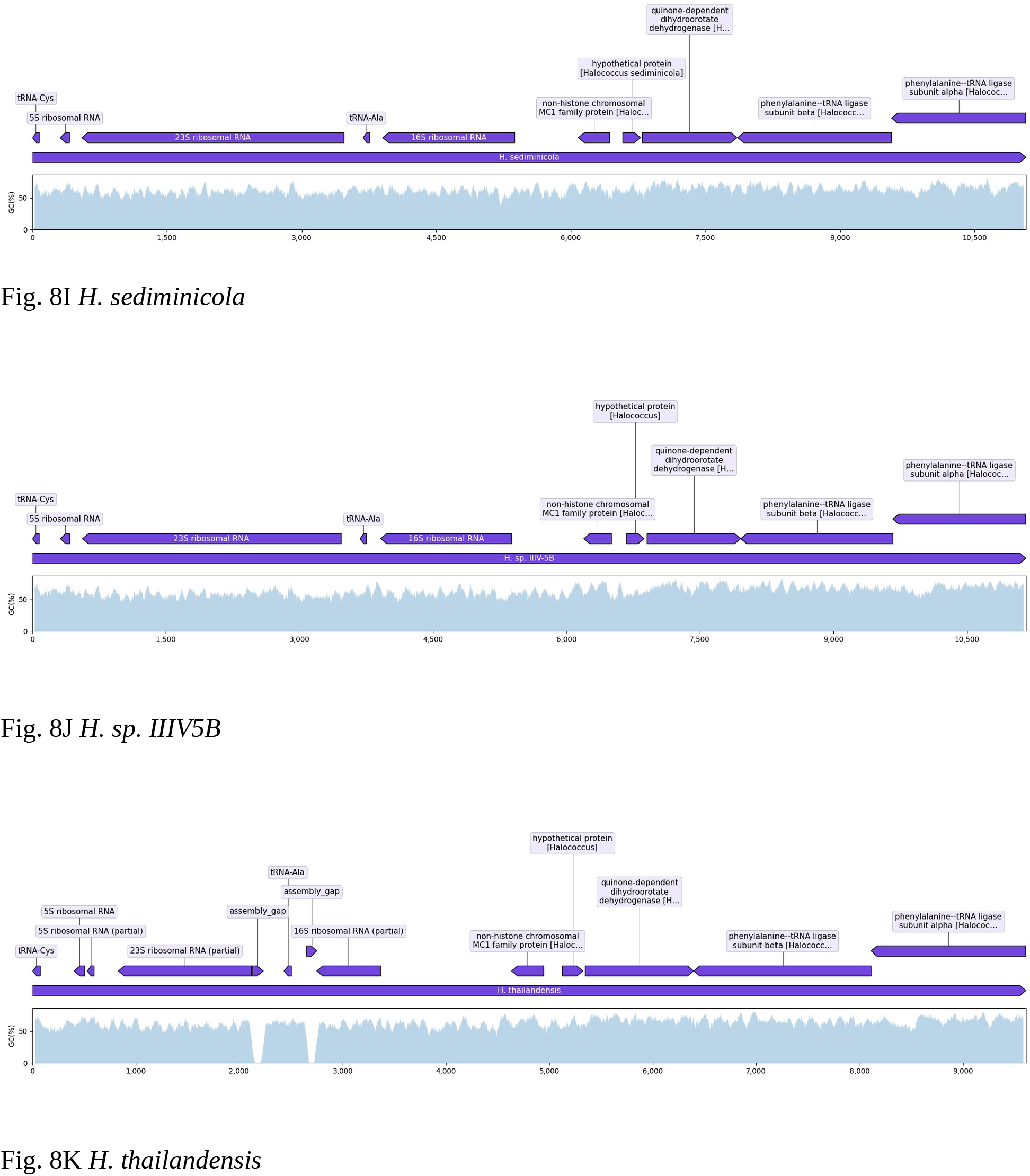
Visualization of the conserved rRNA operon and downstream genes region found across *Halococcus* genomes. The conserved motif downstream of the rRNA operons consists of non-histone chromosomal MC1 family protein, followed by quinone-dependent dihydroorotate dehydrogenase and phenylalanine—tRNA ligase subunit beta and alpha.

While the rRNA operon on the *H. dombrowskii* main chromosome conforms to the known organizational pattern among the Euryarchaeota, the two extra rRNA operons found on plasmid2 (Fig. 9A) and plasmid4 (Fig. 9B) show a markedly different layout, like an inverted sequence of the main rRNA operon, a missing tRNA-Cys site, and a different series of genes downstream of the operons. The downstream genes of the two extrachromosomally encoded rRNA operons are endonuclease NucS, TATA box-binding protein, HtH domain-containing protein, DUF6516 and methyltransferase domain containing protein, as shown below.

**Figure 9.**
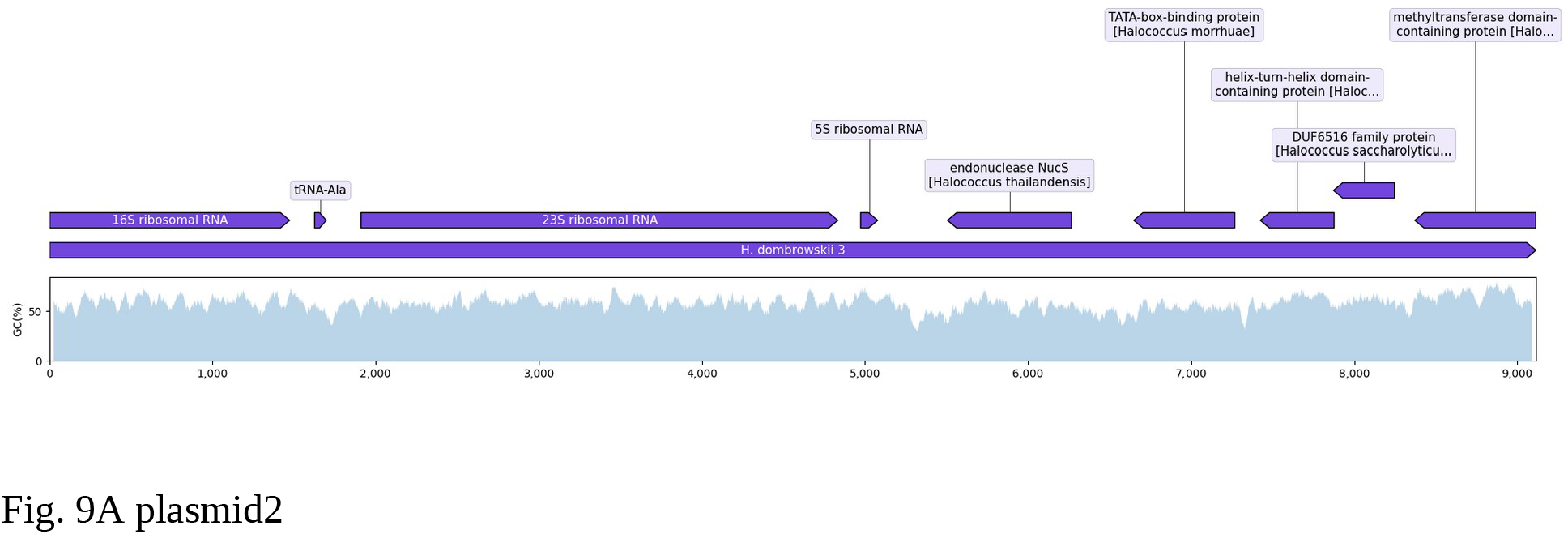
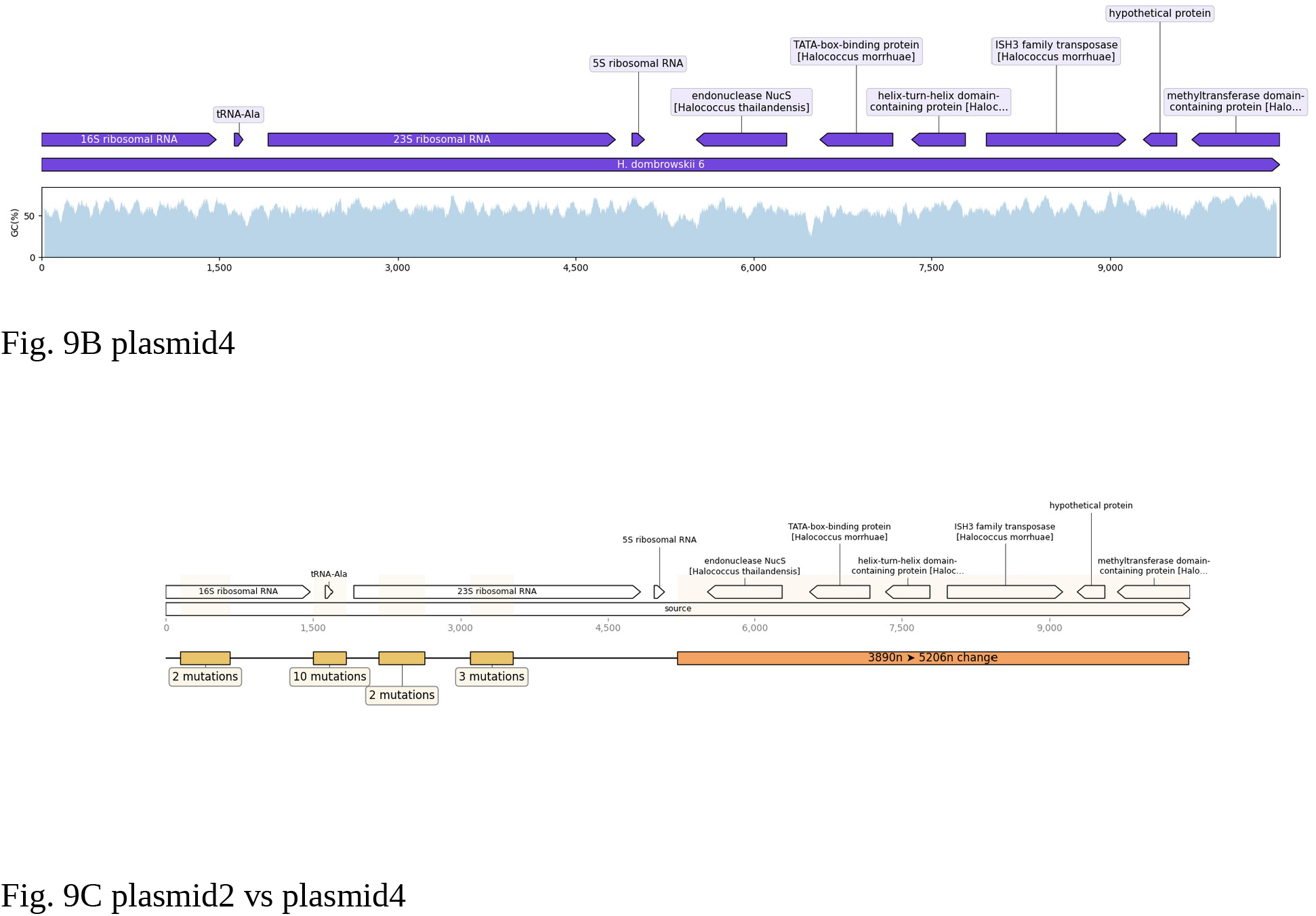
Visualization of the plasmid2 (Fig. 9A) and plasmid4 (Fig. 9B) rRNA operon and downstream genes, showing significant difference from the motifs observed in chromosomal *Halococcus* rRNA operon and downstream gene arrangements.

rRNA operon downstream genes between plasmid2 and plasmid4 exhibits some notable differences between them as well (Fig. 9C). Plasmid4’s rRNA operon downstream region shows an ISH3 family transposase immediately downstream of the HtH domain-containing protein.

Closer inspection suggests that the hypothetical protein following the transposase site is a broken piece of the location matching DUF6516 present on plasmid2’s rRNA operon downstream, as one can see by aligning both protein sequences together (Fig. 10).

**Figure 10.**
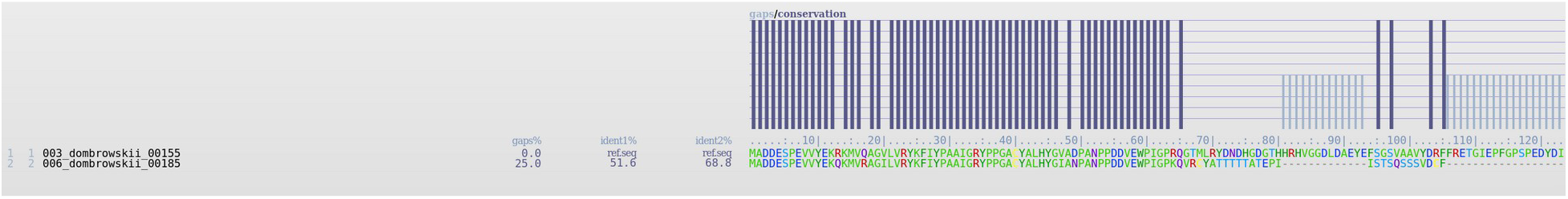
plasmid2’s DUF protein sequence (003) aligned against plasmid4’s hypothetical protein downstream of ISH3 family transposase (006). The plasmid2 DUF6516 protein aligns almost perfectly against the hypothetical protein on plasmid4, until the last half where the plasmid4 hypothetical protein is adjacent to the ISH3 family transposase.

The difference between *H. dombrowskii*’s chromosomal rRNA operon and its plasmid2 and plasmid4 encoded rRNA operons reach beyond just the conserved downstream genes. Alignment of their internal transcribed spacer (ITS) sequences along with the ITS of other *Halococcus* rRNA operons shows an almost one-to-one match between organization of the whole genome *Halococcus* phylogenetic tree and a cladogram derived from just the ITS alignment, indicative of a high degree of conservation that tracks with evolutionary trajectory of the organism. However, there is one notable caveat; *H. dombrowskii*’s chromosomal ITS sequence is more like its phylogenetic neighbor, *H. morrhuae* than the ITS of plasmid2 and plasmid4 (Fig. 11). The difference is enough to place plasmid2 and plasmid4 ITS sequences on a separate clade from the ITS of the host’s own chromosome when aligned together.

**Figure 11.**
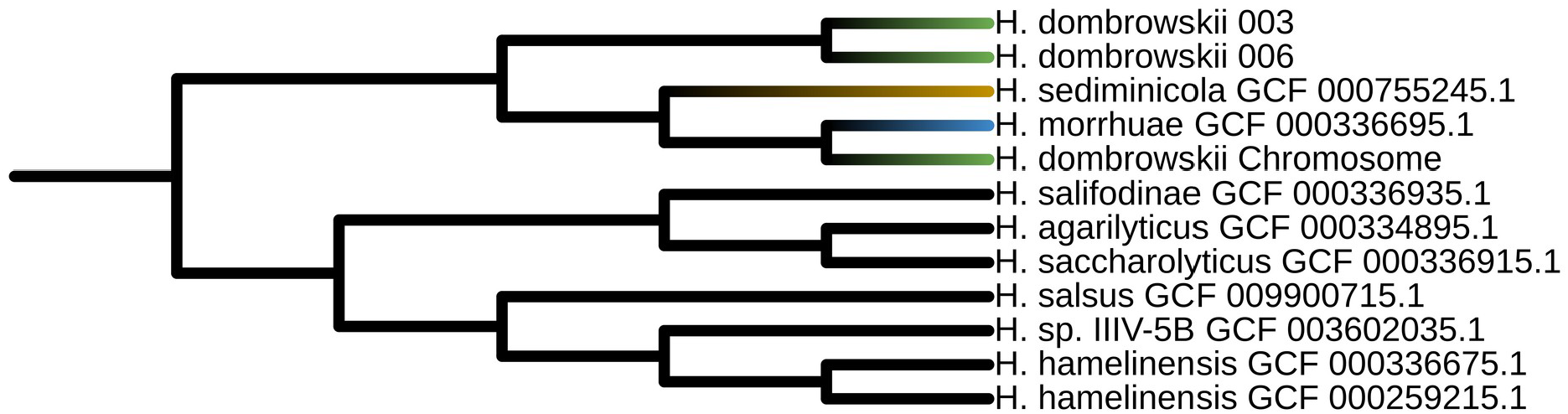
*Halococcus* ITS alignment cladogram. *H. dombrowskii* 003 is ITS region from plasmid2 and *H. dombrowskii* 006 is ITS region from plasmid4. The clade with H. dombrowskii chromosomal ITS shows similarity to its closest neighbors resembling results of CCT genome comparison and whole genome phylogenetic trees. However, the plasmid borne ITS regions (003 and 006) are located on a separate clade. Note that *H. thailandensis* is absent due to gaps in its rRNA and ITS regions.

To verify the independent identity of the three *H. dombrowskii* rRNA operons and conserved downstream genes, we filtered our raw Nanopore data with a minimum 10,000 base pair length cutoff and aligned them against each of the operon and downstream gene blocks, and then sorted only reads unique to each of the operon and downstream genes regions. Visual representation of read length and quality distribution for each of the unique read groups is depicted here (Fig.12A-12C)

**Figure 12.**
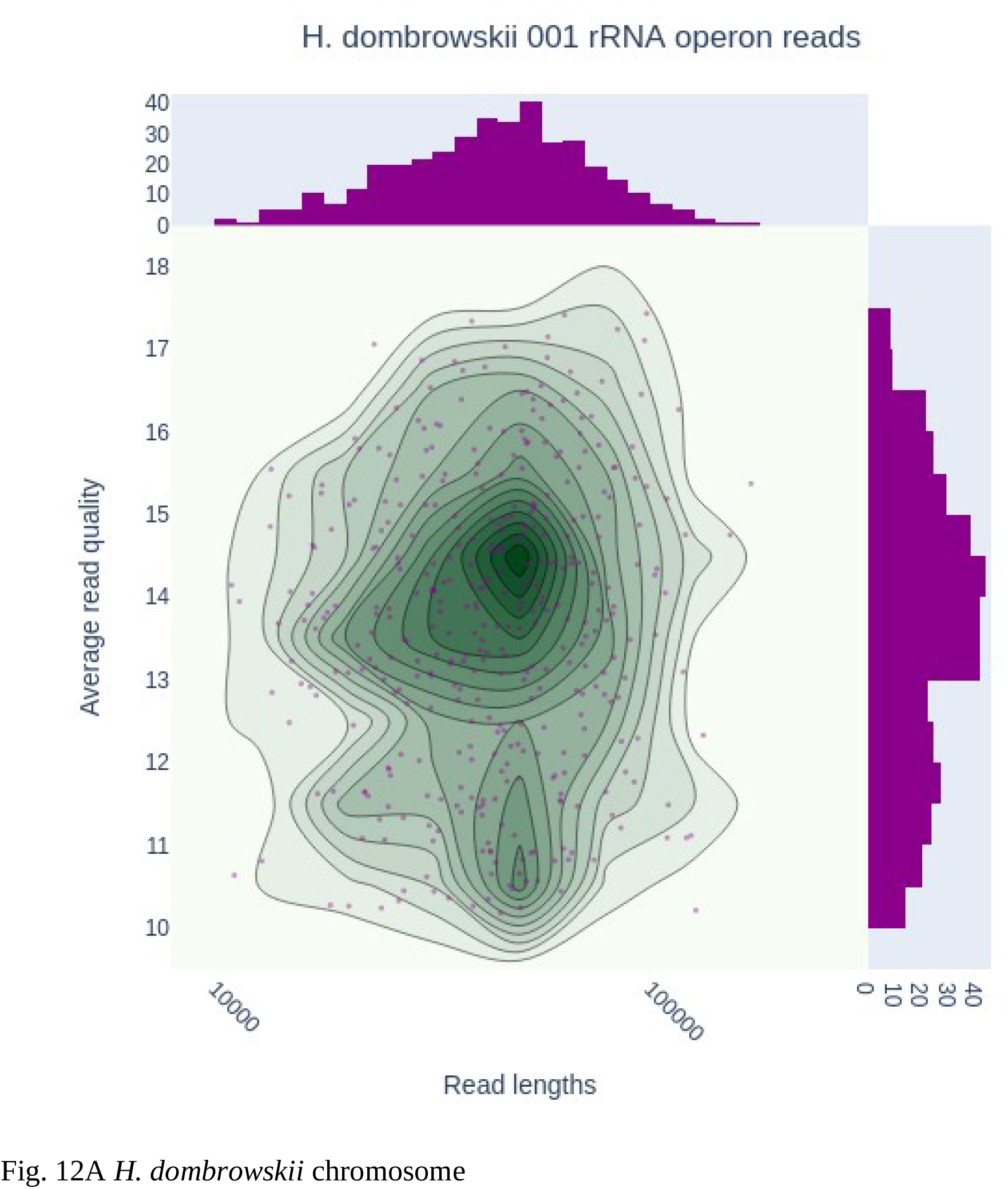
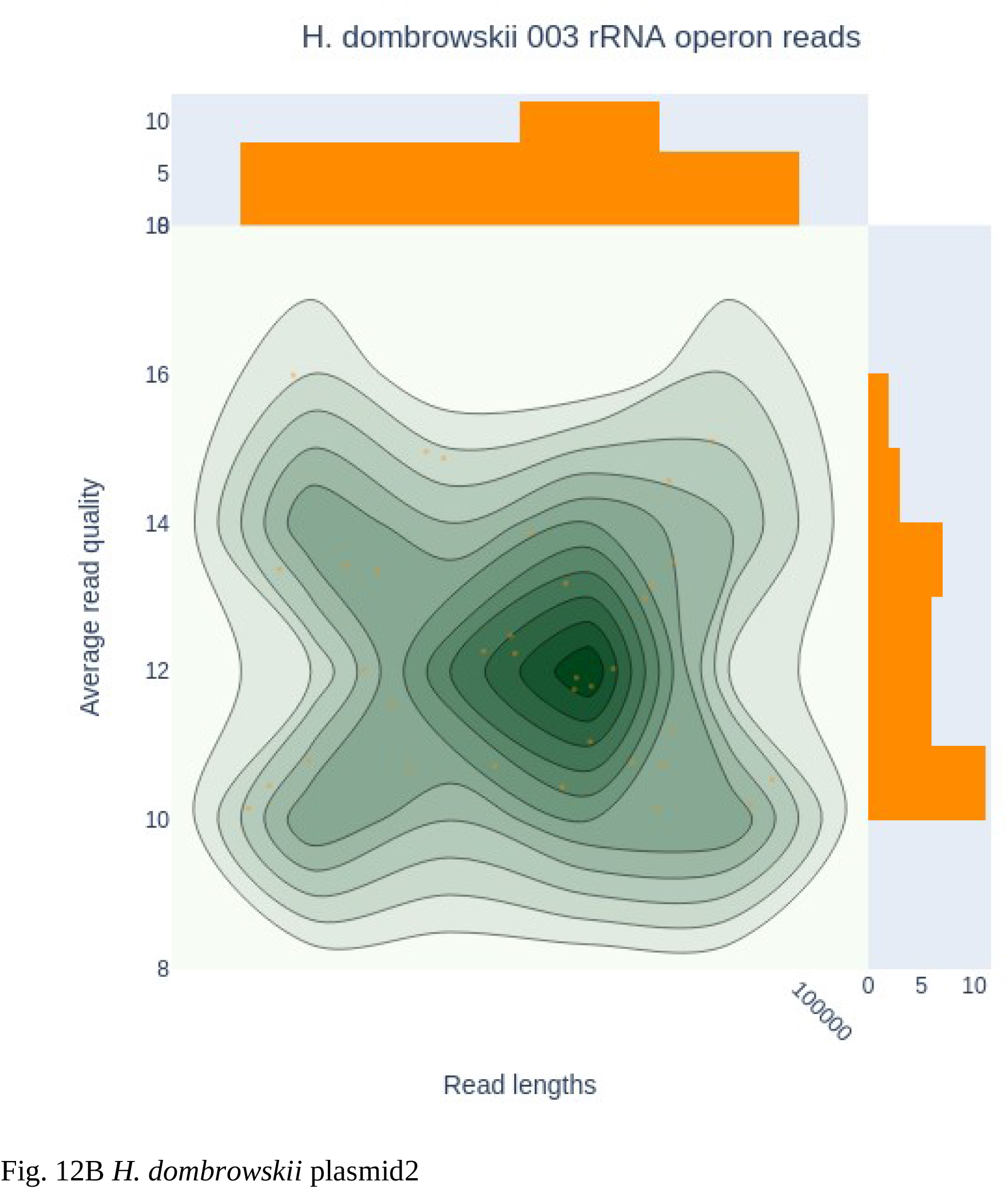
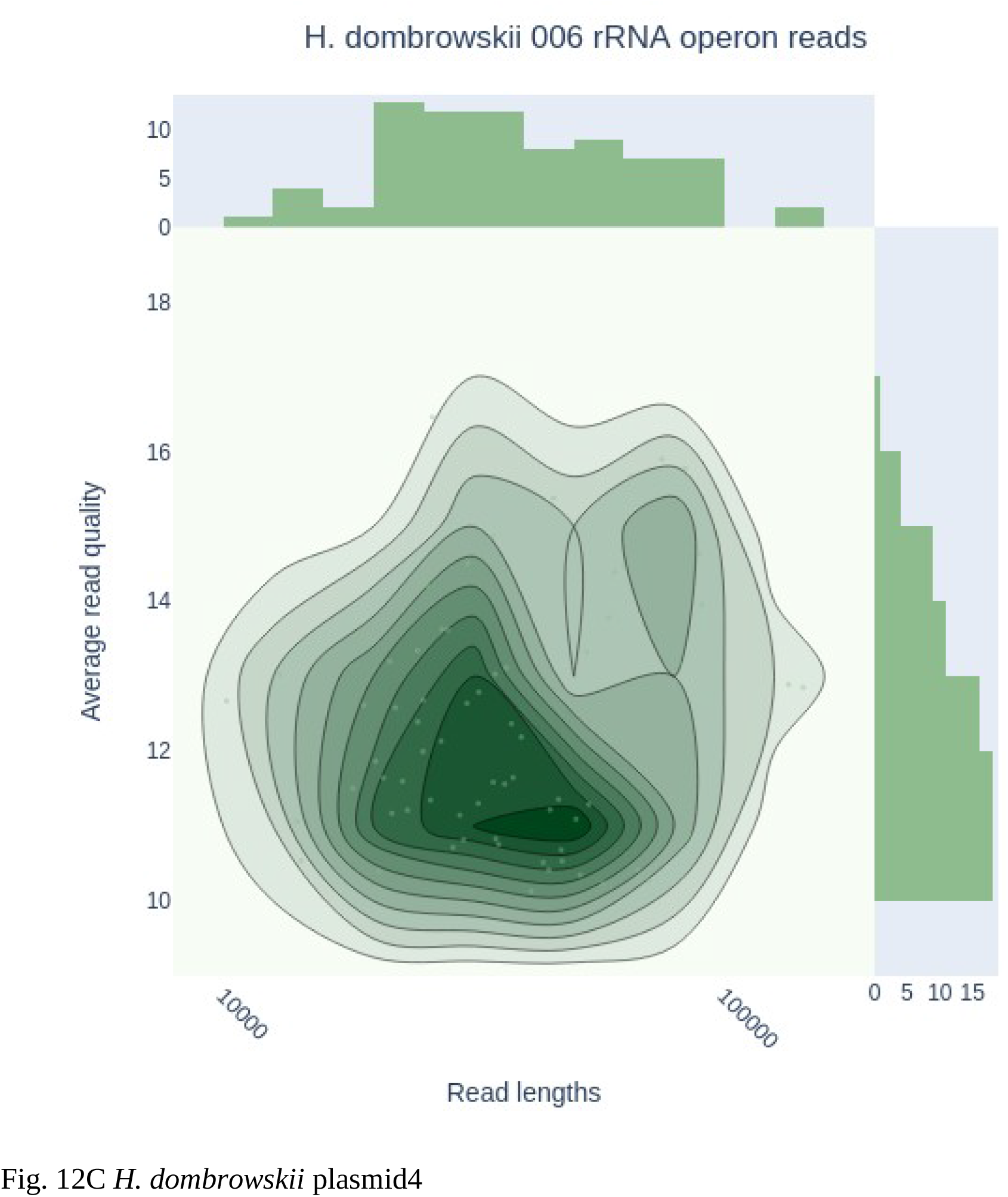
Read quality and length plots of unique reads mapped to *H. dombrowskii* rRNA operon and downstream genes regions. Chromosomal region (Fig. 12A), plasmid2 region (Fig. 12B), and plasmid4 region (Fig. 12C).

The alignment resulted in a total of 384 reads with median read length of 44,512 base pairs mapped uniquely to the chromosomal rRNA operon region and its downstream genes discussed in this paper (Fig. 13A). A total of 35 reads with median read length of 45,812 base pairs map uniquely to the plasmid2 rRNA operon region and its downstream genes (Fig. 13B), and 77 reads with median read length of 35,470 base pairs map uniquely to the plasmid4 rRNA operon region and its downstream genes, including the transposase site unique to the plasmid at near 8000 bp region (Fig. 13C).

**Figure 13.**
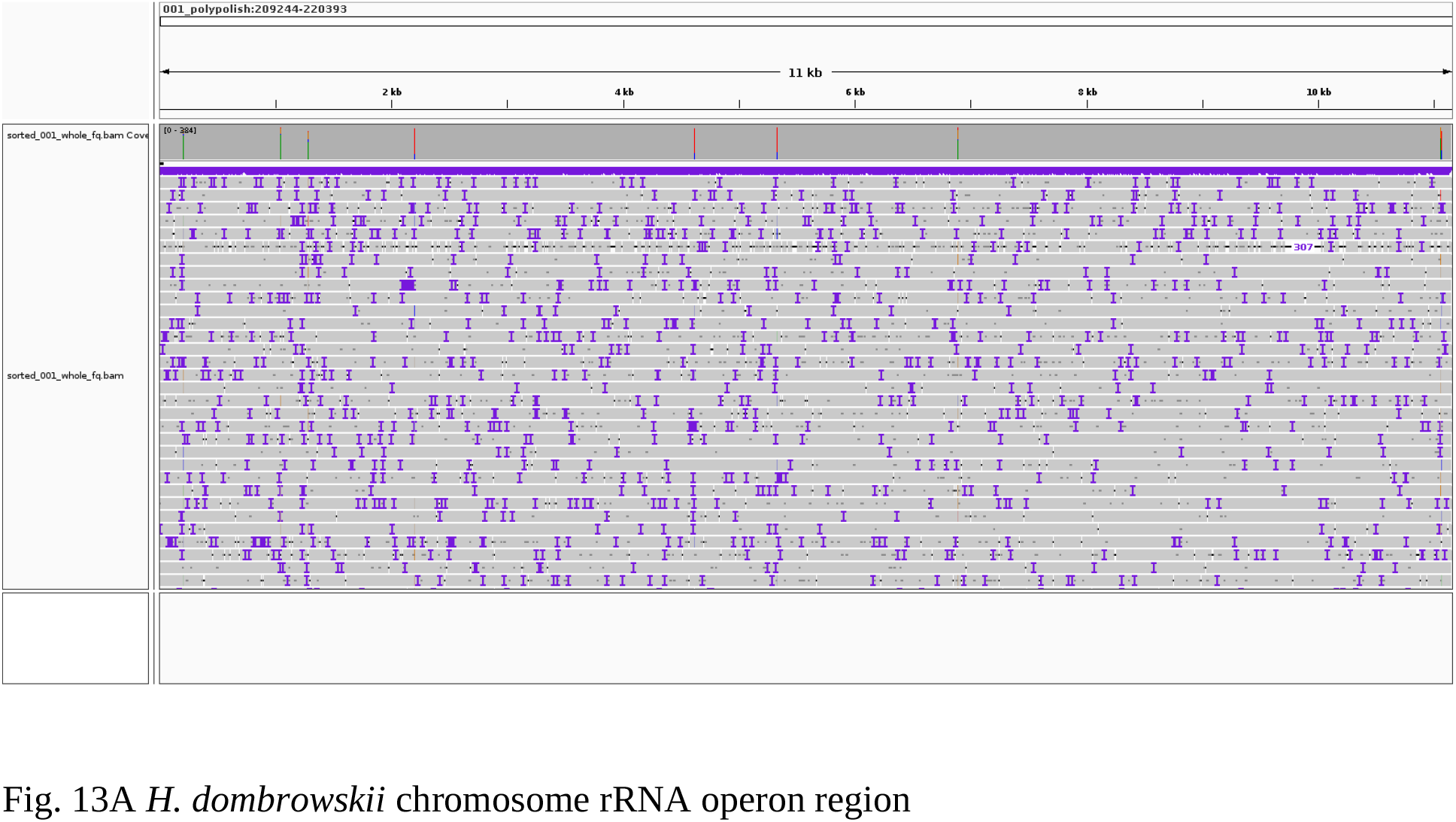
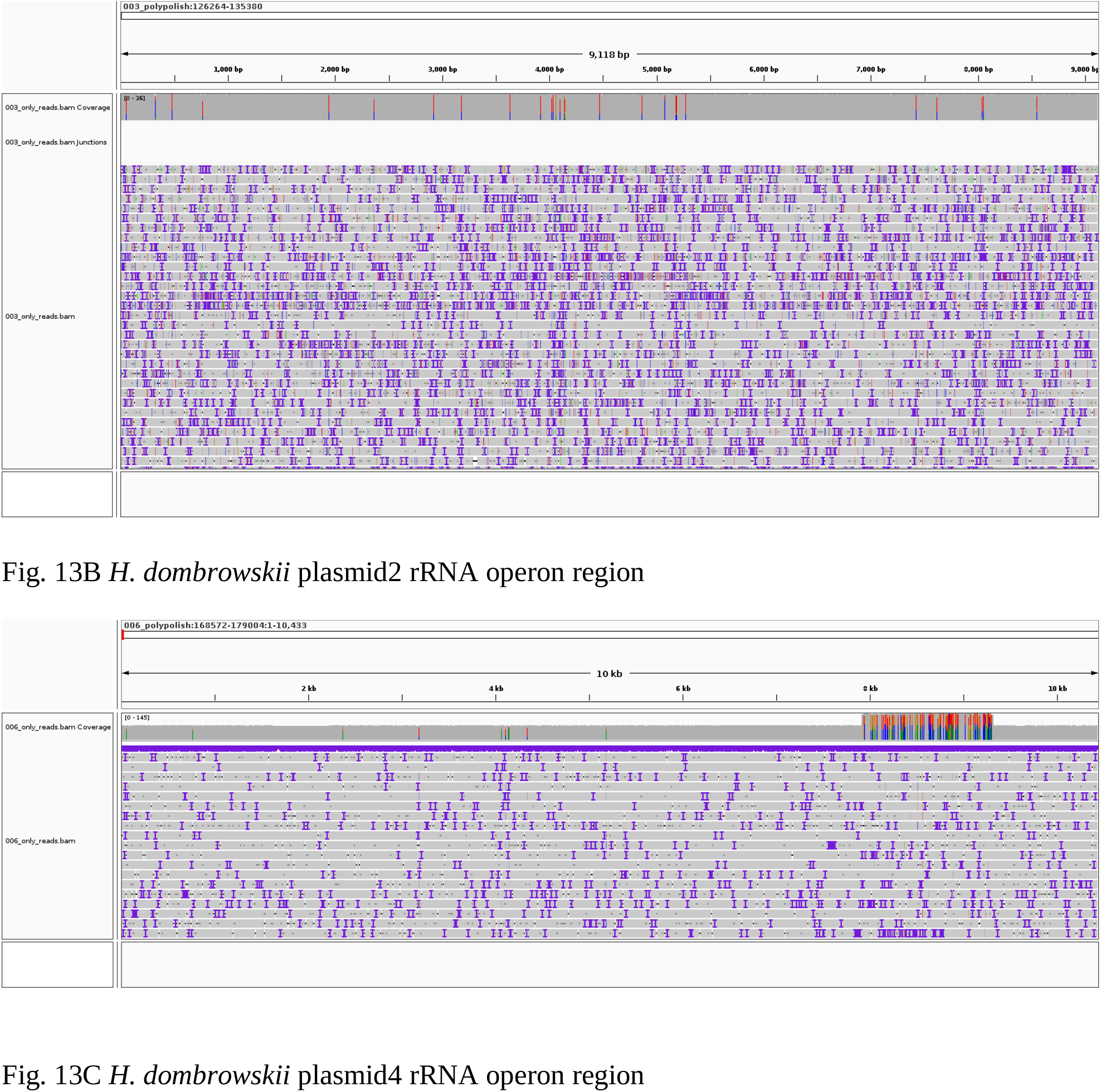
IGV rendering of *H. dombrowskii* reads against chromosomal and plasmid rRNA operon and downstream regions. (13A) the 11150 base pair chromosomal rRNA operon and downstream region; (13B) the 9118 base pair plasmid2 rRNA operon and downstream region; (13C) the 10433 base pair plasmid4 rRNA operon and downstream region, note the high rate of noise matching the ISH3 family transposase site.

The first available *Halococcus dombrowskii* genome presented here depicts a fascinating new portrait of the *Halococcus* species through formal depiction of extrachromosomally encoded rRNA operons with divergent ITS identities. We suspect using long-read platforms to re-sequence and assemble other *Halococcus* genomes could reveal similar features across the genus and support our findings.

The possibility of trackable rRNA operons on extrachromosomal elements and their accompanying sequence elements is especially exciting in light of the still ambivalent origin of Haloarchaea from subsurface salt deposits. Further study on the evolutionary rate of differently located rRNA elements and their relation to 16S-23S ITS divergence could help us create a better molecular clock to shed light on their specific origins.

## DATA AVAILABILITY

The complete genome sequence of *Halococcus dombrowskii* ATCC BAA-364^T^ has been deposited at DDBJ/ENA/GenBank under accession number <u>CP095005</u> for the main chromosome, with five plasmid sequences deposited under accessions <u>CP095006</u>, <u>CP095007</u>, <u>CP095008</u>, <u>CP095009</u>, and <u>CP095010</u>. The raw data from BioProject <u>PRJNA822675</u> were submitted to the NCBI Sequence Read Archive (SRA) and are linked under the BioProject accession above.

## ACKNOWLEDGMENTS

Sequencing and bioinformatics analysis were undertaken at the Hubbard Center for Genome Studies at UNH, supported by NH-INBRE, with the assistance of Kelley Thomas, Joseph Sevigny, and Stephen Simpson. This work was a project of the Microbiology Education through Genome Annotation-New Hampshire (MEGA-NH) program.

## REFERENCES

1. Garrity GM, Holt JG. 2001. Phylum AII. Euryarchaeota., p. 294–300. In Bergey’s manual of systematic bacteriology, 2nd ed. New York : Springer.

2. Farlow WG. 1880. Chapter XLIV. On the nature of the peculiar reddening of salted codfish during the summer time (as observed more particularly at Gloucester, Mass., during the summer of 1878), p. 969–974. In Report of the Commissioner - United States Commission of Fish and Fisheries.

3. Schoop, G. 1935. Halococcus litoralis, ein obligat halphiler Farbstoffbildner. Dtsch Tierärztl Wochenschr 43:817–820.

4. Kocur M, Hodgkiss W. 1973. Taxonomic Status of the Genus Halococcus Schoop. Int J Syst Evol Microbiol 23:151–156.

5. Grant WD. 2001. Genus IV. Halococcus Schoop 1935a, 817AL, p. 311–314. In Bergey’s manual of systematic bacteriology, 2nd ed. New York : Springer.

6. 1989. Approved Lists of Bacterial Names (Amended). ASM Press, Washington (DC). http://www.ncbi.nlm.nih.gov/books/NBK814/. Retrieved 29 July 2022.

7. Montero CG, Ventosa A, Rodriguez-Valera F, Kates M, Moldoveanu N, Ruiz-Berraquero F. 1989. Halococcus saccharolyticus sp. nov., a New Species of Extremely Halophilic Non-alkaliphilic Cocci. Syst Appl Microbiol 12:167–171.

8. Denner EBM, McGenity TJ, Busse H-J, Grant WD, Wanner G, Stan-Lotter H. 1994. Halococcus salifodinae sp. nov., an Archaeal Isolate from an Austrian Salt Mine | Microbiology Society. Int J Syst Evol Microbiol 44:774–780.

9. Stan-Lotter H, Pfaffenhuemer M, Legat A, Busse H-J, Radax C, Gruber C. 2002. Halococcus dombrowskii sp. nov., an archaeal isolate from a Permian alpine salt deposit. Int J Syst Evol Microbiol 52:1807–1814.

10. Goh F, Leuko S, Allen MA, Bowman JP, Kamekura M, Neilan BA, Burns BP. 2006. Halococcus hamelinensis sp. nov., a novel halophilic archaeon isolated from stromatolites in Shark Bay, Australia. Int J Syst Evol Microbiol 56:1323–1329.

11. Wang Q, Li W, Yang H, Liu Y, Cao H, Dornmayr-Pfaffenhuemer M, Stan-Lotter H, Guo G. 2007. Halococcus qingdaonensis sp. nov., a halophilic archaeon isolated from a crude sea-salt sample. Int J Syst Evol Microbiol 57:600–604.

12. Namwong S, Tanasupawat S, Visessanguan W, Kudo T, Itoh T. 2007. Halococcus thailandensis sp. nov., from fish sauce in Thailand. Int J Syst Evol Microbiol 57:2199–2203.

13. Yim KJ, Cha I-T, Whon TW, Lee H-W, Song HS, Kim K-N, Nam Y-D, Lee S-J, Bae J-W, Rhee S-K, Choi J-S, Seo M-J, Roh SW, Kim D. 2014. Halococcus sediminicola sp. nov., an extremely halophilic archaeon isolated from a marine sediment. Antonie Van Leeuwenhoek 105:73–79.

14. Minegishi H, Echigo A, Shimane Y, Kamekura M, Itoh T, Ohkuma M, Usami R. 2015. Halococcus agarilyticus sp. nov., an agar-degrading haloarchaeon isolated from commercial salt. Int J Syst Evol Microbiol 65:1634–1639.

15. Chen S, Sun S, Xu Y, Liu H-C. 2018. Halococcus salsus sp. nov., a novel halophilic archaeon isolated from rock salt. Int J Syst Evol Microbiol 68:3754–3759.

16. Zayas-Rivera J, Rivera-Lopez Y, Velázquez-Méndez M, Romero-Oliveras N, Montalvo-Rodríguez R. 2020. Draft Genome Sequence of a Novel Species of Halococcus (Strain IIIV-5B), an Endophytic Archaeon Isolated from the Leaf Tissue of Avicennia germinans. Microbiol Resour Announc 9:e00421–20.

17. Brito-Echeverría J, López-López A, Yarza P, Antón J, Rosselló-Móra R. 2009. Occurrence of Halococcus spp. in the nostrils salt glands of the seabird Calonectris diomedea. Extrem Life Extreme Cond 13:557–565.

18. Burns BP, Gudhka RK, Neilan BA. 2012. Genome sequence of the halophilic archaeon Halococcus hamelinensis. J Bacteriol 194:2100–2101.

19. Becker EA, Seitzer PM, Tritt A, Larsen D, Krusor M, Yao AI, Wu D, Madern D, Eisen JA, Darling AE, Facciotti MT. 2014. Phylogenetically driven sequencing of extremely halophilic archaea reveals strategies for static and dynamic osmo-response. PLoS Genet 10:e1004784.

20. Yim KJ, Kim B-Y, Lee H-W, Song HS, Nam Y-D, Choi J-S, Choi H-J, Seo M-J, Yoon C, Kim K-N, Kim D, Rhee J-K, Roh SW. 2014. Draft genome sequence of the extremely halophilic archaeon Halococcus sediminicola CBA1101T isolated from a marine sediment sample. Mar Genomics 18PB:145–146.

21. Selim S, Hagagy N. 2016. Genome sequence of carboxylesterase, carboxylase and xylose isomerase producing alkaliphilic haloarchaeon Haloterrigena turkmenica WANU15. Genomics Data 7:70–72.

22. Dombrowski H. 1963. Bacteria from Paleozoic Salt Deposits. Ann N Y Acad Sci 108:453–460.

23. McGenity TJ, Gemmell RT, Grant WD, Stan-Lotter H. 2000. Origins of halophilic microorganisms in ancient salt deposits. Environ Microbiol 2:243–250.

24. Wick RR, Judd LM, Holt KE. 2019. Performance of neural network basecalling tools for Oxford Nanopore sequencing. Genome Biol 20:129.

25. De Coster W, D’Hert S, Schultz DT, Cruts M, Van Broeckhoven C. 2018. NanoPack: visualizing and processing long-read sequencing data. Bioinforma Oxf Engl 34:2666–2669.

26. Kolmogorov M, Yuan J, Lin Y, Pevzner PA. 2019. Assembly of long, error-prone reads using repeat graphs. 5. Nat Biotechnol 37:540–546.

27. Chen S, Zhou Y, Chen Y, Gu J. 2018. fastp: an ultra-fast all-in-one FASTQ preprocessor. Bioinforma Oxf Engl 34:i884–i890.

28. Wick RR, Judd LM, Cerdeira LT, Hawkey J, Méric G, Vezina B, Wyres KL, Holt KE. 2021. Trycycler: consensus long-read assemblies for bacterial genomes. Genome Biol 22:266.

29. Li H. 2016. Minimap and miniasm: fast mapping and de novo assembly for noisy long sequences | Bioinformatics | Oxford Academic. Bioinformatics 32:2103–2110.

30. Wick RR, Holt KE. 2019. Benchmarking of long-read assemblers for prokaryote whole genome sequencing. F1000Research 8:2138.

31. Vaser R, Šikić M. 2021. Time-and memory-efficient genome assembly with Raven. 5. Nat Comput Sci 1:332–336.

32. Shafin K, Pesout T, Lorig-Roach R, Haukness M, Olsen HE, Bosworth C, Armstrong J, Tigyi K, Maurer N, Koren S, Sedlazeck FJ, Marschall T, Mayes S, Costa V, Zook JM, Liu KJ, Kilburn D, Sorensen M, Munson KM, Vollger MR, Monlong J, Garrison E, Eichler EE, Salama S, Haussler D, Green RE, Akeson M, Phillippy A, Miga KH, Carnevali P, Jain M, Paten B. 2020. Nanopore sequencing and the Shasta toolkit enable efficient de novo assembly of eleven human genomes. Nat Biotechnol 38:1044–1053.

33. Seemann T. 2014. Prokka: rapid prokaryotic genome annotation. Bioinforma Oxf Engl 30:2068–2069.

34. Lee MD. 2019. GToTree: a user-friendly workflow for phylogenomics. Bioinforma Oxf Engl 35:4162–4164.

35. Lee MD. 2019. Applications and Considerations of GToTree: A User-Friendly Workflow for Phylogenomics. Evol Bioinforma Online 15:1176934319862245.

36. Letunic I, Bork P. 2021. Interactive Tree Of Life (iTOL) v5: an online tool for phylogenetic tree display and annotation. Nucleic Acids Res 49:W293–W296.

37. Li H, Durbin R. 2010. Fast and accurate long-read alignment with Burrows-Wheeler transform Bioinforma Oxf Engl 26:589–595.

38. Vaser R, Sovic I, Nagarajan N, Sikic M. 2017. Fast and accurate de novo genome assembly from long uncorrected reads. Genome Res gr.214270.116.

39. Wick RR, Holt KE. 2022. Polypolish: Short-read polishing of long-read bacterial genome assemblies PLOS Comput Biol 18:e1009802.

40. Simão FA, Waterhouse RM, Ioannidis P, Kriventseva EV, Zdobnov EM. 2015. BUSCO: assessing genome assembly and annotation completeness with single-copy orthologs. Bioinforma Oxf Engl 31:3210–3212.

41. Manni M, Berkeley MR, Seppey M, Zdobnov EM. 2021. BUSCO: Assessing Genomic Data Quality and Beyond. Curr Protoc 1:e323.

42. Manni M, Berkeley MR, Seppey M, Simão FA, Zdobnov EM. 2021. BUSCO Update: Novel and Streamlined Workflows along with Broader and Deeper Phylogenetic Coverage for Scoring of Eukaryotic, Prokaryotic, and Viral Genomes. Mol Biol Evol 38:4647–4654.

43. Grant JR, Arantes AS, Stothard P. 2012. Comparing thousands of circular genomes using the CGView Comparison Tool. BMC Genomics 13:202.

44. Li H, Handsaker B, Wysoker A, Fennell T, Ruan J, Homer N, Marth G, Abecasis G, Durbin R, 1000 Genome Project Data Processing Subgroup. 2009. The Sequence Alignment/Map format and SAMtools. Bioinforma Oxf Engl 25:2078–2079.

45. Sievers F, Wilm A, Dineen D, Gibson TJ, Karplus K, Li W, Lopez R, McWilliam H, Remmert M, Söding J, Thompson JD, Higgins DG. 2011. Fast, scalable generation of high-quality protein multiple sequence alignments using Clustal Omega. Mol Syst Biol 7:539.

46. Quinlan AR, Hall IM. 2010. BEDTools: a flexible suite of utilities for comparing genomic features Bioinforma Oxf Engl 26:841–842.

47. Thorvaldsdóttir H, Robinson JT, Mesirov JP. 2013. Integrative Genomics Viewer (IGV): high-performance genomics data visualization and exploration. Brief Bioinform 14:178–192.

48. Galperin MY, Makarova KS, Wolf YI, Koonin EV. 2015. Expanded microbial genome coverage and improved protein family annotation in the COG database. Nucleic Acids Res 43:D261–D269.

49. Yip WSV, Vincent NG, Baserga SJ. 2013. Ribonucleoproteins in Archaeal Pre-rRNA Processing and Modification. Archaea 2013:e614735.

