## Supplemental for "Analysis of the complete genome sequence for *Halococcus dombrowskii* ATCC BAA-364^T^"

### Supplemental Materials

#### SUPPLEMENTAL METHODS

##### Characterizing and filtering raw MinION and Illumina reads

All MinION reads the M2 and M3 long-read samples were characterized using NanoPlot v.1.39.0, using the following commands:

NanoPlot -o m2_all_plot --summary sequencing_summary.txt --loglength --readtype 1D --plots kde --color blueviolet --title "SUP basecalled M2 reads"

NanoPlot -o m3_all_plot --summary sequencing_summary.txt --loglength --readtype 1D --plots kde --color crimson --title "SUP basecalled M3 reads"

We then used FiltLong v.0.2.1 on only the pass quality reads from M2 and M3 (minimum Q score of 10) to generate working data for each group, screening for minimum read length of 1000bp and then keeping the top scoring 90% of the output from each group:

filtlong ../../m2/m2_pass.fastq.gz --min_length 1000 --keep_percent 90 > m2_1k_90.fastq

filtlong ../../m3/m3_pass.fastq.gz --min_length 1000 --keep_percent 90 > m3_1k_90.fastq

Resulting working read sets were characterized with NanoPlot again:

NanoPlot -o m2_1k_90 --fastq m2_1k_90.fastq --loglength --readtype 1D --plots kde --color blueviolet --title "m2 1000bp cutoff 90%"

NanoPlot -o m3_1k_90 --fastq m3_1k_90.fastq --loglength --readtype 1D --plots kde --color crimson --title "m3 1000bp cutoff 90%"

We observed fragmented (high contig numbers) Flye-based exploratory genome assemblies based solely on M2/M3 data. To address the possibility that we're working with highly degraded DNA, we also combined the M2 and M3 passed reads set into a single set prior to the filtering process described above. The combined reads were filtered separately as a third data group referred to as M23.

filtlong --min_length 1000 --keep_percent 90 ../../m23/m23_pass.fastq.gz > m23_1k_90.fastq

NanoPlot -o m23_1k_90 --fastq m23_1k_90.fastq --loglength --readtype 1D --plots kde --color magenta --title "m23 1000bp cutoff 90%"

Fastp v.0.23.1 was used to filter short reads from Illumina HiSeq using the command below:

fastp -i HdomA3_CGCTCATT-TATAGCCT_L001_R1_001.fastq -o filtered_HdomA3_R1.fastq -I HdomA3_CGCTCATT-TATAGCCT_L001_R2_001.fastq -O filtered_HdomA3_R2.fastq --

unpaired1 unpaired1.fastq --unpaired2 unpaired2.fastq --detect_adapter_for_pe --correction --overrepresentation_analysis

**Genome assembly**

Trycycler v.0.5.3 provided the backend for Halococccus dombrowskii genome assembly, based on output of four different assemblers; Flye v.2.9-b1768, Miniasm v.0.3-r179 & Minipolish v0.1.3, Raven v.1.7.0, and finally Shasta v.0.8.0. M2, M3 and M23 filtered read sets were used to create 16 subsampled reads each. The subsampled reads corresponded to 4 assemblies per 4 different target assemblers, producing 48 assemblies in total.

Subsampling command for Trycycler was as follows:

```
trycycler subsample --count 16 --reads m2_1k_90.fastq --out_dir m2_1k_90
trycycler subsample --count 16 --reads m3_1k_90.fastq --out_dir m3_1k_90
trycycler subsample --count 16 --reads m23_1k_90.fastq --out_dir m23_1k_90
```

A bash script was used to generate assemblies from each of the subsampled reads (Sup #), and resulting assemblies were annotated with Prokka v.1.14.6 to check for rRNA and CRISPR spacer site counts. Prokka command was as follows:

```
for F in *.fasta; do
N=$(basename $F .fasta);
prokka --kingdom Archaea --rawproduct --locustag $N --outdir $N --prefix $N $F;
done
```

GToTree v.1.6.31 was used to generate cladograms for each of the M2/M3/M23 set based assembly groups using the command below, and annotation file was curated manually to show rRNA and CRISPR count per assembly on itol.embl.de visualization platform.

```
GToTree -f fasta_files.txt -H Archaea -m genome_to_id_map.tsv -o m2_1k_90_tree
 -j 8

GToTree -f fasta_files.txt -H Archaea -m genome_to_id_map.tsv -o m3_1k_90_tree
 -j 8

GToTree -f fasta_files.txt -H Archaea -m genome_to_id_map.tsv -o m23_1k_90_tre
e -j 8
```

18 assemblies were chosen out of 48 based on rRNA count and CRISPR spacer site count and pooled into a separate directory to create clusters of raw reads around contigs. We used the filtered M23 reads for this step.

```
trycycler cluster --assemblies assemblies/*.fasta --reads m23_1k_90.fastq --ou
t_dir trycycler --threads 20
```

The clustering process generated 39 clusters containing 165 contigs in total. Reconciliation of the contigs using below command filtered the clusters down to a final 6, clusters 1, 2, 3, 4, 6 and 9. Note that the below command needs to be run on a per-cluster basis.

trycycler reconcile --reads m23_1k_90.fastq --cluster_dir trycycler/cluster_*

Multiple sequence alignment was performed on each of the remaining clusters via below command:

trycycler msa --cluster_dir trycycler/cluster_*

The filtered M23 reads were partitioned to each of the aligned cluster sequences:

trycycler partition --reads m23_1k_90.fastq --cluster_dirs trycycler/cluster_* --threads 20

Final consensus sequence was generated on per contig basis, and the contigs were combined into final raw assembly using below command:

trycycler consensus --cluster_dir trycycler/cluster_*

cat trycycler/cluster_*/7_final_consensus.fasta > trycycler/consensus.fasta

Composition of the final unpolished assembly is as follows:

```
######################################################
-------------------------------------------------
 755405 A
 743911 T
1229564 C
1236343 G
-------------------------------------------------
Total gapped sequence length is: 3965223
-------------------------------------------------
Total ungapped sequence length is: 3965223
-------------------------------------------------
GC content in consensus.fasta is 62.18 %
######################################################
```

The raw assembly was polished using an in-house script (Sup #) using bwa v.0.7.17-r1198-dirty, Racon v.1.4.20, Medaka v.1.5.0 and Polypolish v.0.5.0.

polish_everything.sh m23_1k_90.fastq filtered_HdomA3_R1.fastq.gz filtered_HdomA3_R2.fastq.gz

Composition of the final polished assembly is as follows:

```
######################################################
```

-------------------------------------------------
 755483 A
 744011 T
1229572 C
1236400 G
-------------------------------------------------
Total gapped sequence length is: 3965466
-------------------------------------------------
Total ungapped sequence length is: 3965466
-------------------------------------------------
GC content in H_dombrowskii.fasta is 62.18 %
##################################################

**Characterizing and annotating the genome**

We performed additional QC using BUSCO v.5.3.2 and archaea_odb10 profile using the command below:

busco -m genome -i H_dombrowskii.fasta -o H_dombrowskii_busco -l archaea_odb10

GToTree was used again to generate a whole genome phylogenetic tree of Halococcus dombrowskii against other Halococcus genomes in NCBI. Halococcus reference genomes were queried on NCBI Genbank using below parameters, which returned 10 genome accessions.

txid2249[Organism:exp] AND ("latest refseq"[filter] AND all[filter] NOT anomal
ous[filter])

GCF_000336675.1 Halococcus hamelinensis 100A6
GCF_000259215.1 Halococcus hamelinensis 100A6 (a distinct genome from the sample above)
GCF_000336915.1 Halococcus saccharolyticus DSM 5350
GCF_000336935.1 Halococcus salifodinae DSM 8989
GCF_000336695.1 Halococcus morrhuae DSM 1307
GCF_000336715.1 Halococcus thailandensis JCM 13552
GCF_009900715.1 Halococcus salsus ZJ1
GCF_000755245.1 Halococcus sediminicola CBA1101
GCF_000334895.1 Halococcus agarilyticus 197A
GCF_003602035.1 Halococcus sp. IIIV-5B

Phylogenetic tree was built using both our local Halococcus dombrowskii genome and the NCBI Halococcus genome accession using below command:

GToTree -a 1103_midroot_halococcus_genus_gcf.txt -f fasta_files.txt -H Archaea -m genome_to_id_map.tsv -o midroot_halococcus_tree -j 8

In preparation for a more detailed Prokka annotation, , we curated a list of known genomes from class Halobacteria on NCBI using below parameters, returning 501 genome accessions:

txid183963[Organism:exp] AND ((latest[filter] OR "latest refseq"[filter]) AND all[filter] NOT anomalous[filter])

NCBI-genome-download v.0.3.1 was used with Halobacteria genome accessions from above to compile a Halobacterial protein sequences library:

```
ncbi-genome-download archaea -A 1103_halobacteria_class_gcf.txt -F protein-fasta -o halobacteria_class_gcf_protein
```

We then split the assembly into contigs and annotated them using Prokka and the protein sequences library.

```
for F in *.fasta; do
N=$(basename $F .fasta);
prokka --kingdom Archaea --protein halobacteria_gcf.faa --rawproduct --genus Halococcus --locustag $N --outdir $N --prefix $N $F;
done
```

**Visualization and analysis *Halococcus* spp. genomes**

A docker instance of CGview Comparison Tool was used to compare the Halococcus dombrowskii against other Halococcus species on whole genome level. NCBI-genome-download script was used again to obtain genbank files of the 10 Halococcus genomes:

```
ncbi-genome-download archaea -A 1103_halococcus_genus_gcf.txt -F genbank -o halococcus_gcf_genbank
```

We initialized CGview Comparison Tool with our Halococcus dombrowskii genbank file using below command:

```
sudo docker run --rm -v "$(pwd)":/dir -u "$(id -u)":"$(id -g)" -w /dir pstothard/cgview_comparison_tool build_blast_atlas.sh -i H_dombrowskii.gbk
```

Other Halococcus species genbank files were placed in comparison_genomes folder of the resulting directory, and analysis initialized with following command:

```
sudo docker run --rm -v "$(pwd)":/dir -u "$(id -u)":"$(id -g)" -w /dir pstothard/cgview_comparison_tool build_blast_atlas.sh -p H_dombrowskii -z large --custom "title='Halococcus dombrowskii genome vs other Halococcus species' titleFontSize=150 global_label=F legend=T legendFontSize=30 gc_skew=T gene_labels=F details=F gcSkewColorNeg=blue gcSkewColorPos=red"
```

Once analysis and figure generation was complete, we created a zoomed in view of the rRNA regions of the Halococcus genomes using below command:

```
sudo docker run --rm -v "$(pwd)":/dir -u "$(id -u)":"$(id -g)" -w /dir
pstothard/cgview_comparison_tool create_zoomed_maps.sh -p H_dombrowskii -c 212000 -z 35
```

rRNA regions and downstream genes were manually isolated using samtools v.1.14 with both Prokka-generated and NCBI genbank files as reference. The DNA sequences around 10,000 bases in length were re-annotated with Prokka and used with Geneblocks v.1.2.2 script to generate figures used in this paper.

The same process was used to isolate the Internal Transcribed Spacer (ITS) sequences from the Halococcus genomes, and Clustal Omega, hosted on the website of the EBI, was used to align them and generate the cladogram used for the figure.

**Isolating reads unique to *H. dombrowskii* rRNA regions**

M23 raw reads were filtered again with a minimum length of 10,000 bp to cut down on spurious mapping to only parts of our regions of interest. Top 90% of reads at least 10,000 bp length were isolated using the below filtlong v.0.2.1 (R.R. Wick) command:

```
filtlong --min_length 10000 --keep_percent 90 m23_pass.fastq.gz > m23_pass_90_10k.fastq
```

Three sequence blocks containing rRNA operon and downstream genes were manually isolated from Halococcus dombrowskii genome with samtools and Prokka-annotated genbank file: the 11150 bp chromosomal (contig 001) region (NZ_CP095005.1) between 209244-220393, the 9117 bp plasmid 2 (contig 003) region (NZ_CP095007.1) between 126264-135380, and the 10433bp plasmid 4 (contig 006) region (NZ_CP095009.1) between 168572-179004. Minimap v.2.24-r1122 and Bedtools v.2.29.1 (26) were used to isolate reads mapping uniquely to the rRNA operon and downstream gene regions on either H. dombrowskii's chromosomal rRNA, plasmid 2 or plasmid 4 using shell commands outlined below:

The chromosomal sequence block was processed first:

```
samtools faidx dombrowskii_001_rnaOperon.fasta
minimap2 -ax map-ont dombrowskii_001_rnaOperon.fasta ../m23_pass_90_10k.fastq | samtools
view -bhS | samtools sort -o 001_reads.bam
samtools index 001_reads.bam
```

A bed file was written covering the whole of the chromosomal rRNA operon and genes region:

```
001_polypolish:209244-220393    1       11150
```

And Bedtools intersect was used to isolate reads that encompass the region described in the BED file:

```
bedtools intersect -wa -a 001_reads.bam -b 001_whole.bed -F 1.0 | samtools sort -o
sorted_001_whole_fq.bam
samtools index sorted_001_whole_fq.bam
```

```
bedtools bamtofastq -i 001_whole_fq.bam -fq 001_whole_fq.fastq
bedtools bamtobed -i 001_reads.bam > 001_reads.bed
bedtools intersect -wa -a 001_reads.bed -b 001_whole.bed -F 1.0 > 001_whole_fq.bed
awk '{print $4}' 001_whole_fq.bed | sort > 001_cov_reads.txt
```

Plasmid 2 sequence block was processed next:

```
minimap2 -ax map-ont dombrowskii_003_rnaOperon.fasta ../m23_pass_90_10k.fastq
| samtools view -bhS | samtools sort -o 003_reads.bam
samtools index 003_reads.bam
```

Another bed file was written covering the whole of plasmid 2 rRNA operon and genes region:

```
003_polypolish:126264-135380   1       9117
```

```
bedtools intersect -wa -a 003_reads.bam -b 003_whole.bed -F 1.0 | samtools sort -o
003_whole_fq.bam
samtools index 003_whole_fq.bam
bedtools bamtofastq -i 003_whole_fq.bam -fq 003_whole_fq.fastq
awk 'END{print NR/4}' 003_whole_fq.fastq
bedtools bamtobed -i 003_whole_fq.bam > 003_whole_fq.bed
awk '{print $4}' 003_whole_fq.bed | sort > 003_cov_reads.txt
```

Followed by plasmid 4 sequence block:

```
minimap2 -ax map-ont dombrowskii_006_rnaOperon.fasta ../m23_pass_90_10k.fastq | samtools
view -bhS | samtools sort -o 006_reads.bam
samtools index 006_reads.bam
```

Final bed file was written covering the whole of plasmid 4 rRNA operon and genes region:

```
006_polypolish:168572-179004   1       10433
```

```
bedtools intersect -wa -a 006_reads.bam -b 006_whole.bed -F 1.0 | samtools sort -o
006_whole_fq.bam
samtools index 006_whole_fq.bam
bedtools bamtofastq -i 006_whole_fq.bam -fq 006_whole_fq.fastq
bedtools bamtobed -i 006_whole_fq.bam > 006_whole_fq.bed
awk '{print $4}' 006_whole_fq.bed | sort > 006_cov_reads.txt
```

We used grep to compare and isolate unique fastq headers from reads mapped to each of the sequence blocks. Below command returns 77 reads unique to the plasmid 4 sequence block:

```
grep -Fxv -f 003_cov_reads.txt ../006_operon/006_cov_reads.txt | wc -l
```

And below command returns 35 reads unique to plasmid 2 sequence block:

```
grep -Fxv -f ../006_operon/006_cov_reads.txt 003_cov_reads.txt | wc -l
```

The unique read headers are sorted and combined to create a list we can compare against reads aligned to the chromosomal sequence block:

```
cat 003_cov_reads.txt ../006_operon/006_cov_reads.txt | sort | uniq > 003_006_reads.txt
```

Running below command with above read header list returns 384 reads unique to the chromosomal sequence block:

```
grep -Fxv -f 003_006_reads.txt ../001_operon/001_cov_reads.txt | wc -l
```

[Seqtk]() v.1.3-r117-dirty is used to create fastq files containing only reads unique to each of the sequence blocks.

For reads mapping uniquely to rRNA operon and downstream regions on plasmid 2:

```
seqtk subseq 003_whole_fq.fastq  003_only_reads.txt > 003_only_reads.fastq
```

And for reads mapping uniquely to rRNA operon and downstream regions on plasmid 4:

```
seqtk subseq 006_whole_fq.fastq  006_only_reads.txt > 006_only_reads.fastq
```

NanoPlot is used to visualize and characterize the unique reads fastq files:

```
NanoPlot -o 001_whole_fq_plot --fastq 001_whole_fq.fastq --loglength --readtype 1D --plots kde --color darkmagenta --title "H. dombrowskii 001 rRNA operon reads"
```

```
NanoPlot -o 003_only_reads_plot --fastq 003_only_reads.fastq --loglength --readtype 1D --plots kde --color darkorange --title “H. dombrowskii 003 rRNA operon reads”
```

```
NanoPlot -o 006_only_reads_plot --fastq 006_only_reads.fastq -loglength --readtype 1D --plots kde --color darkseagreen --title “H. dombrowskii 006 rRNA operon reads”
```

IGV was used to show mapped reads against the rRNA operon and downstream genes regions.
